# Iron-dependent mutualism between *Chlorella sorokiniana* and *Ralstonia pickettii* forms the basis for a sustainable bioremediation system

**DOI:** 10.1101/2021.06.15.446916

**Authors:** Deepak Rawat, Udita Sharma, Pankaj Poria, Arran Finlan, Brenda Parker, Radhey Shyam Sharma, Vandana Mishra

**Author notes:** **Corresponding Authors:** 1. Prof. Vandana Mishra, Department of Environmental Studies, Bioresources & Environmental Biotechnology Laboratory, University of Delhi, Delhi-110007, INDIA. 2. Dr Brenda Parker, Department of Biochemical Engineering, Bernard Katz Building, University College London, Gower Street, London WC1E 6BT, UK, 02076799789. 3. Prof. Radhey Shyam Sharma, Department of Environmental Studies, Bioresources & Environmental Biotechnology Laboratory, University of Delhi, Delhi-110007, INDIA.

## Abstract

Mutualism between microalgae and bacteria is ubiquitous, but remains underexplored as a basis for biodegradation of anthropogenic pollutants. In industrial systems, poor iron uptake by microalgae limits growth, bioprocessing efficacy, and bioremediation potential. Iron supplementation is costly and ineffective because iron remains insoluble in aqueous medium and biologically unavailable. In aquatic environments, microalgae develop an association with bacteria that solubilize iron by production of siderophore, which increases the bioavailability of iron as a public good. Algae, in exchange, provides dissolved organic matter to bacteria to sustain such interkingdom associations. Therefore, using a case study of azo dye degradation, we combine environmental isolations and synthetic ecology as a workflow, establishing a microbial community to degrade industrially relevant Acid Black 1 dye. We create a mutualism between previously non-associated chlorophyte alga *Chlorella sorokiniana* and siderophore-producing bacterium *Ralstonia pickettii*, based on the eco-evolutionary principle of exchange of iron and carbon. This siderophore-mediated increased iron bioavailability increases reductive iron uptake, growth rate, and azoreductase-mediated dye degradation of microalga. In exchange, *C. sorokiniana* produces galactose, glucose, and mannose as major extracellular monosaccharides, supporting bacterial growth. We propose a mechanism whereby extracellular ferrireductase is crucial for azoreductase-mediated dye degradation in microalgae. Our work demonstrates that bioavailability of iron, which is often overlooked in industrial bio-designs, governs microalgal growth and enzymatic processes. Our results suggest that algal-bacterial consortia based on the active association are a self-sustainable mechanism to overcome existing challenges of micronutrient availability in bioremediation systems and their industrial translation.

## Introduction

Microalgae have potential in the creation of a circular economy, preventing pollution, and enabling the reuse of water as a resource. The textile industry releases ~5 million tons of untreated wastewater annually into waterways, accounting for 1/5^th^ of the global water pollution and releasing ecotoxic compounds like aromatic amines arising from incomplete degradation of dye by environmental microbes [1]. Exposure to dye contaminants arising from textile industries disproportionately affects emerging economies like China, India, and Bangladesh, hubs of textile dyeing, where environmental regulations are relaxed, and wastewater treatment and management are inefficient [2, 3]. Therefore, there is an urgent need to address the sustainability of this sector.

Commonly used physico-chemical methods aim for only color removal and recently employed zero-liquid discharge techniques convert liquid waste into untreatable hazardous sludge [4]. Though alternative biological methods can be advantageous for industrial use in developing countries, they have lower operational costs, generate less sludge, and result in degradation of dye toxicants into environmentally benign compounds [5], yet their potential remains unrealized, owing to constraints related to biomass upscaling under unfavorable environmental conditions of wastewater and integration in the current infrastructure.

Microalgae such as *Chlamydomonas, Chlorella, Dunaliella, Micratinium, Scenedesmus*, and *Phaeodactylum* have been investigated for their use in wastewater treatment and in biorefinery processes to recover valuable products of interest [6–11]. In particular, species of *Chlorella* and *Scenedesmus* have received attention for wastewater applications, including textile, due to their rapid doubling times and tolerance for a wide range of nutrient conditions [8, 12, 13]. However, in industrial wastewater, algal processes may be challenged by extrinsic constraints of micronutrient availability and intrinsic constraints of micronutrient utilization. The inability of algae to take up complexed iron limits its efficacy to treat industrial wastewater [14], which has low iron concentrations and even low bioavailability due to high alkalinity (pH 8.5-10) [15, 16]. Microalgae require Fe^2+^ for photosynthesis, respiration, nitrogen-fixation, uptake of nutrients, and metabolism of reactive oxygen species [17]. Although abundant in the environment, iron remains unavailable to algae due to its presence as insoluble ferric (Fe^3+^) or oxyhydroxides or as bound to minerals and organics. In alkaline conditions, like in wastewater from the textile industry, iron precipitates out or dissolute slowly [16, 18]. Therefore, bioavailable iron remains an insignificant fraction in aqueous environments, thus, acts as a major limiting factor for upscaling of microalgal-based processing of wastewater treatment.

Phototrophs and heterotrophs occupy distinct ecological niches but may engage in mutualism to complement their physiological capabilities and metabolic versatilities to enhance fitness [19–23]. Photoautotrophic microalgae represent a source of dissolved organic carbon that may be supplied to heterotrophic bacteria. In contrast to microalgae, many species of heterotrophic bacteria produce siderophores, low molecular weight iron chelators, as a part of the highly efficient iron-uptake mechanism in iron-stressed environments [21]. To overcome their physiological limitations, autotrophic algae exchange dissolved organic matter with heterotrophic bacteria in lieu of non-bioavailable micronutrients and other metabolites [24, 25].

In bioremediation designs and bioprocesses, single-species cultures have several physiological limitations that challenge their use in industrial wastewater treatment [26]. For example, microalgal monocultures are challenged with limited bioavailability of micronutrients (like Fe, Mn, vitamins), the metabolic burden of maintaining essential cellular processes under stressful conditions [27, 28]. In textile wastewater remediation, bacterial monocultures are challenged by the sensitivity of azoreductase enzymes for oxygen [29], necessitating a complex two-stage approach for complete dye removal. They require an oxygen-limiting environment to reduce azo dyes to aromatic amines extracellularly. Further treatment requires a well-oxygenated environment for oxidative degradation of aromatic amines [4]. Photosynthetic algae have been reported to degrade dye [13]; however, azoreductases in microalgae have not been studied previously. Unlike bacteria, microalgae could regulate external oxygen concentrations, therefore, could provide a simpler single-stage solution for complete azo dye degradation [12]. Also, algal-mediate oxygen release in wastewater facilitates bacterial growth in primary treatment processes and accelerates the degradation of organic pollutants [30], which will complement the degradation of aromatic amines in dye wastewater.

Therefore, employing mutualistic microalgae and bacteria would ensure their use as a sustainable option in remediating toxicants and combating environmental stresses by distributing tasks to share the metabolic burden [31]. In freshwater and marine ecosystems, siderophore-producing bacteria develop a mutualistic association with other microbes, including microalgae, therefore, govern the structure and function of microbial communities [25, 32]. In alkaline conditions, found in textile wastewater, bacterial siderophores, due to chelation and higher dissolution, could enrich iron in the immediate environment to use as public goods [16, 33]. This ensures higher primary productivity of microalgae for a continued exchange of dissolved organic matter. However, the impact of such eco-evolutionary principles of mutualism between organisms in improving the sustainability of bioremediation processes is largely underexplored [26, 34, 35].

Therefore, the present study was aimed to (i) assess the role of siderophore-producer bacteria in enhancing the growth of microalgae under iron-limiting conditions and (ii) to assess the potential of the microalgal-bacterial consortium in degrading azo dyes. We isolated siderophore-producer *Ralstonia pickettii* from industrial wastewater and demonstrated their ability to undergo mutualistic association with previously non-associated freshwater microalgae *Chlorella sorokiniana*, and assessed the efficacy of mutualism-assisted azoreductase-mediated microalgal remediation of textile dyes.

## Results and discussion

### Assessment of siderophore production in bacteria and dye degradation in algae

Out of seven bacterial isolates purified from the untreated textile wastewater, five isolates showed relatively high siderophore production (Supplementary Fig. S1). *Ralstonia pickettii* PW2, *Serratia plymuthica* PW1, and *S. liquefaciens* PW71 grew within 24 h on deferrated CAS agar plates and showed siderophore activity, i.e., the appearance of the orange-yellow zone due to siderophore-mediated removal of Fe from blue-colored CAS-HDTMA-Fe complex [40]. *Stenotrophomonas maltophilia* PW5 and *S. maltophilia* PW6 showed high growth and siderophore production but only after 96 h of incubation. *S. rhizophila* PW3 and *S. rhizophila* PW72, however, failed to grow on CAS agar plates. *S. plymuthica* PW1, *S. liquefaciens* PW71, and *R. pickettii* PW2 produced siderophore in decreasing order of concentration, i.e., 15.26±1.3 > 13.28±0.9 > 10.85±0.7 μMmL^-1^ (Table 1). Arnow’s and Csaky’s assay confirmed a catecholate-type siderophore is produced by *Serratia plymuthica* PW1 (81.10±9.8 μMmL^-1^), *Ralstonia pickettii* PW2 (97.43±16.8 μMmL^-1^), and *Serratia liquefaciens* PW71 (103.1±8.3 μMmL^-1^). On the other hand, hydroxamate-type of siderophore is produced by *S. maltophilia* PW5 (37.86±0.4 μMmL^-1^) and *S. maltophilia* PW6 (17.73±0.2 μMmL^-1^) (Table 1). As *S. rhizophila* PW3 and *S. rhizophila* PW72 showed low culturability in iron-limiting conditions, they were omitted from further experiments.

**Table 1:** Characterization of siderophore production in bacterial strains isolated from textile wastewater.

| Bacteria | Concentration ($\mu\text{M mL}^{-1}$) | | |
| --- | --- | --- | --- |
|  | Universal-CAS assay (w.r.t. Desferrioxamine mesylate) | Catecholate- Arnow's assay (w.r.t. 2'3-Dihydroxybenzoic acid) | Hydroxamate- Csaky's assay (w.r.t. Desferrioxamine mesylate) |
| *Serratia plymuthica* PW1 | 15.26±1.28 | 81.10±9.85 | 1.35±1.03 |
| *Ralstonia pickettii* PW2 | 10.85±0.70 | 97.43±16.86 | NA |
| *Stenotrophomonas rhizophila* PW3 | 1.31±0.40 | NA | NA |
| *Stenotrophomonas maltophilia* PW5 | 1.28±0.34 | 2.77±1.20 | 37.86±0.46 |
| *Stenotrophomonas maltophilia* PW6 | 1.63±0.07 | 2.43±0.33 | 17.73±0.26 |
| *Serratia liquefaciens* PW71 | 13.28±0.92 | 103.1±8.33 | NA |
| *Stenotrophomonas rhizophila* PW72 | 0.91±0.63 | 7.4±6.86 | NA |

Our study also reports *S. plymuthica* PW1 and *S. liquefaciens* PW71 from textile dye effluent produce catecholate siderophores (Table 1), which is common among stress-tolerant bacterial genera [51]. Different *Serratia* species thrive well in contaminated environments such as textile dye effluent and petroleum [52]. *R. pickettii* PW2, a β-proteobacterium, has also been reported to produce siderophores under iron stress[53].

Out of the five algal species screened, the two freshwater microalgae *C. sorokiniana, Scenedesmus* sp., and the freshwater cyanobacterium *O. animalis* were observed to degrade AB 1 dye. After 72 h, the AB1 degradation potential of microalgae was in decreasing order as *C. sorokiniana* (82.10±1.6%) > *Scenedesmus* sp. (31.04±3.1%) > *O. animalis* (30.36±1.8%). The marine algal species *Phaeodactylum tricornutum* 1052/6 and 1055/1 did not decolorize AB1 dye under the conditions tested. *Chlorella* and *Scenedesmus* sp. have also been known to degrade a wide range of pollutants such as textile dyes [12, 54], heavy metals [21], pesticides [6], and aromatic hydrocarbons [54]. Therefore, we selected *Chlorella* and *Scenedesmus* for testing and developing an algal-bacterial consortium for dye degradation.

### Iron and carbon dependent mutualism between *Chlorella sorokiniana* and *Ralstonia pickettii*

The exudates from *C. sorokiniana* and *Scenedesmus* sp. were used as a source of dissolved organic matter for cultivating bacteria and identifying microalgal-bacterial combination having the potential to act as a consortium (Fig. 1E). All five bacterial isolates grow well on filter-sterilized exudate of *C. sorokiniana* as a sole source of carbon. On the contrary, on exudate of *Scenedesmus* sp.*, S. plymuthica* PW1 showed moderate growth in 20 h while the growth of *R. pickettii* PW2 and *S. liquefaciens* PW71 remained insignificant. *S. maltophilia* PW5 and *S. maltophilia* PW6 did not grow on the exudate of *Scenedesmus* sp. (Supplementary Fig. S2B).

**Fig. 1.**
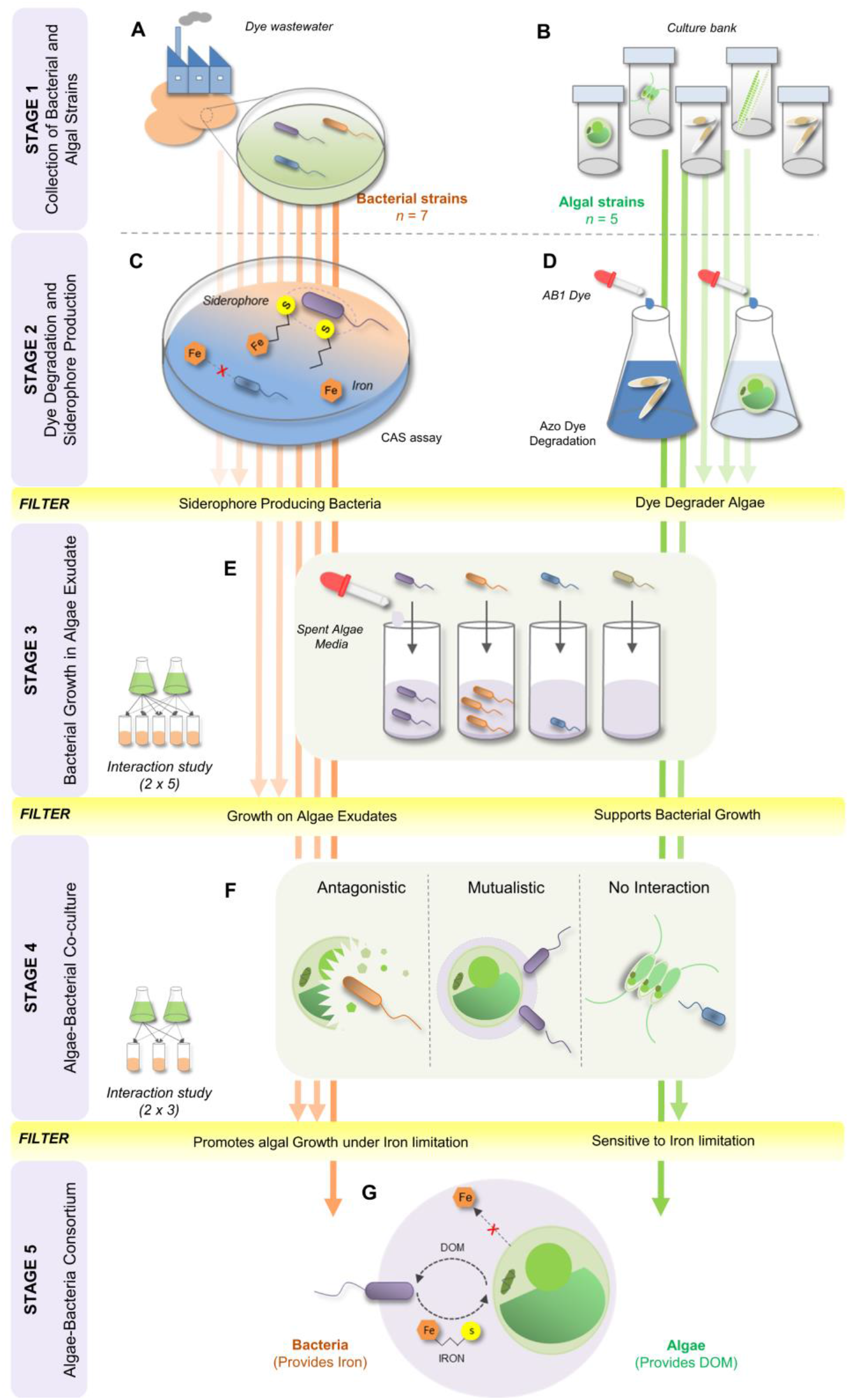
The study design explaining different stages of experiments to identify a consortium of previously non-associated algae and bacteria. The stages include **A** isolation of bacterial strains from textile wastewater collected from Panipat Industrial area, Haryana (India); **B** cultivation of freshwater and marine algal strains; **C** assessment of siderophore production in bacterial strains using Schwyn and Neilands’s universal Chrome Azurol S (CAS) assay; **D** assessment of dye degradation potential of algae strains using Acid Black 1 (AB1) dye; **E** interaction study between siderophore producing bacteria and dye degrader microalgae to identify bacterial strains that could sustain on algae-derived DOM secreted in algal exudates; **F** algal-bacterial co-culturability assessment to study different types of symbiotic associations viz. antagonism, mutualism, or no interaction between the two organisms, and **G** identification of algal-bacterial model consortium based on the active exchange of iron and DOM.

Different combinations of consortia with microalgal (*C. sorokiniana* / *Scenedesmus* sp.) and siderophore-producer bacterial (*S. plymuthica* PW1/ *R. pickettii* PW2/ *S. liquefaciens* PW71) partner were assessed for dye degradation under iron limitation (Fig. 1F). Microalgal cell count in consortium culture was compared with that of axenic microalgal culture to categorize its interaction with a bacterial partner as mutualistic, antagonistic, and neutral. In contrast with axenic microalgal culture (~4×10^6^ cells mL^-1^), *C. sorokiniana* in co-culture with *R. pickettii* PW2 showed a significant increase in cell count at 200 h (~6×10^6^ cells mL^-1^), suggesting a mutualistic association (Fig. 2A). However, *S. plymuthica* PW1 exerted an antagonistic effect on the growth of *C. sorokiniana*, whereas, the interaction between *S. liquefaciens* PW71 and *C. sorokiniana* remained neutral (Fig. 2A). During early growth, the interaction between *Scenedesmus* sp. and *S. plymuthica* PW1 remained neutral; however, later, the interaction turned as antagonistic (Fig. 2A). Whereas the interaction of *Scenedesmus* sp. with both *R. pickettii* PW2 and *S. liquefaciens* PW71 was neutral. *Scenedesmus* showed a higher growth (~12×10^6^ cells mL^-1^) than axenically grown *Chlorella* (~5×10^6^ cells mL^-1^), suggesting an effective iron-uptake mechanism under iron-limiting conditions.

**Fig. 2.**
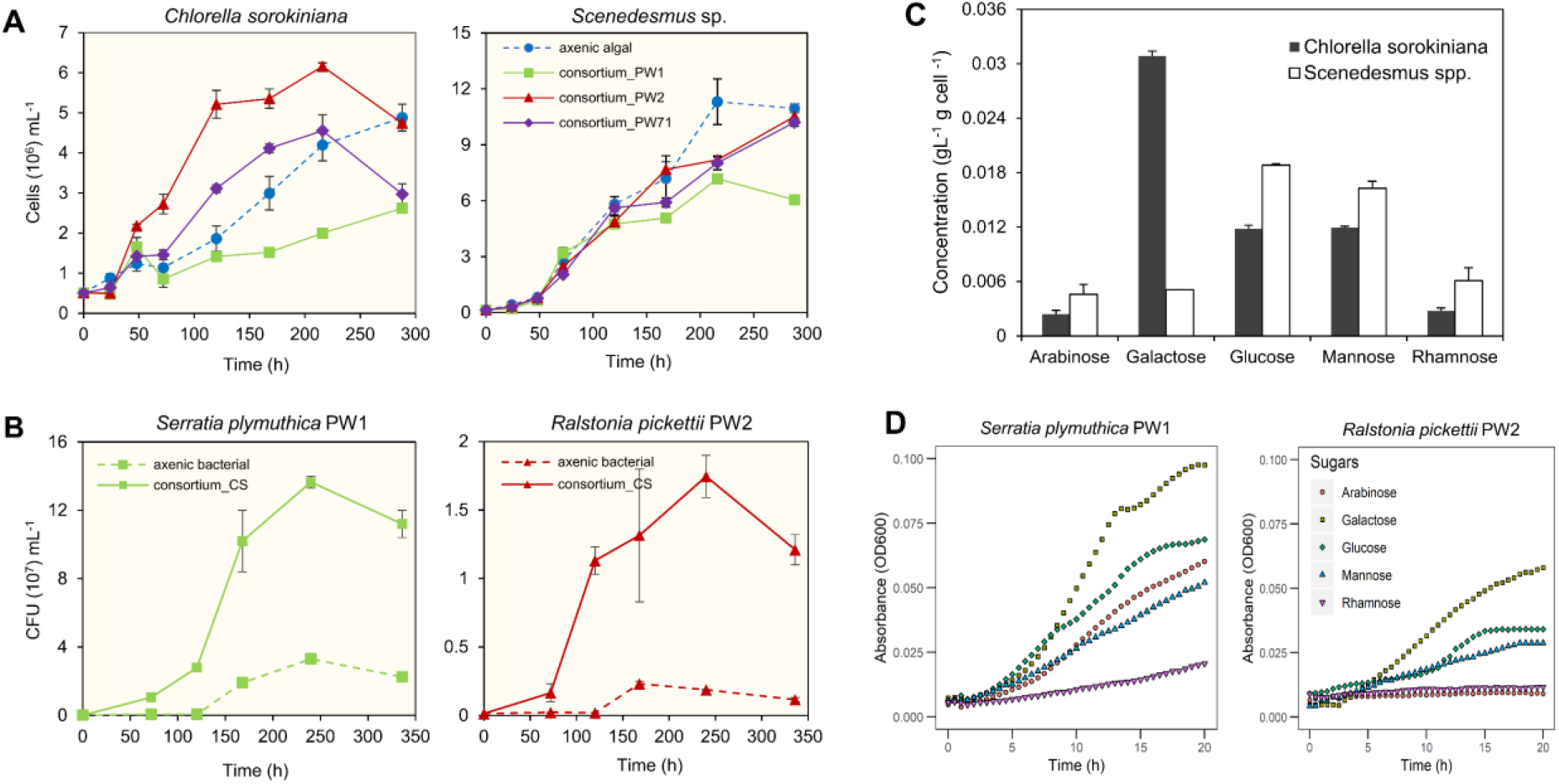
Assessment of algal and bacterial growth in co-culture experiments. **A** Algal cell counts from co-culture experiments suggesting mutualistic effect of *Ralstonia pickettii* PW2 on *C. sorokiniana*. *Serratia plymuthica* PW1 on the other hand showed antagonistic effect on *C. sorokiniana*, and *Serratia liquefaciens* PW71 showed neutral effect. The effect of bacteria on growth of *Scenedesmus* sp. in co-culture was less prominent. **B** The assessment of the CFUs of bacterial strains in algal-bacterial co-culture suggest the growth promoting effect of *Chlorella sorokiniana* on PW1 and PW2. **C** Anion-exchange chromatography suggests a difference in the glycosyl composition in the EPS content of two freshwater microalgae. **D** The bacterium *Serratia plymuthica* PW 1 also showed high culturability in all 5 sugars suggesting the generalist behavior of the strain. Whereas, bacterium *Ralstonia pickettii* PW2 showed limited culturability only in galactose followed by glucose and mannose. Galactose was preferred by both the bacteria.

Growth parameters such as carrying capacity (k), intrinsic growth rate (r), doubling time (Dt), and area under the curve (auc) provided better insights into the population ecology of microalgae [42] (Supplementary Fig. S2, Table S1, and Table S2). As indicated by a steeper slope of log-phase in the growth curve (Fig. 3A), the growth rate (r) of *C. sorokiniana* in consortium with *R. pickettii* PW2 (5.02±1.0×10^-2^ h^-1^) remained significantly higher than that of axenically grown microalgae (1.95±0.3×10^-2^ h^-1^) (*p*=0.000). However, the carrying capacity (k) of *C. sorokiniana* remains unchanged with and without culturing with *R. pickettii* PW2 (co-culture: 4.97±0.1×10^6^ cells vs axenic culture: 4.82±0.4×10^6^ cells; *p*=1.000). In the consortium, *C. sorokiniana* showed a higher population turnover during the early log-phase and reached the stationary phase earlier (at 100 h) than that grown axenically (~270 h) though carrying capacity (k) remained similar. A significantly higher area under curve (auc) in the consortium (11.01±0.4×10^8^) of *R. pickettii* PW2 in comparison with its axenic culture (6.01±0.5×10^8^) (*p*=0.000) indicate mutualism between *C. sorokiniana* and *R. pickettii* PW2 (Fig. 3A).

**Fig. 3.**
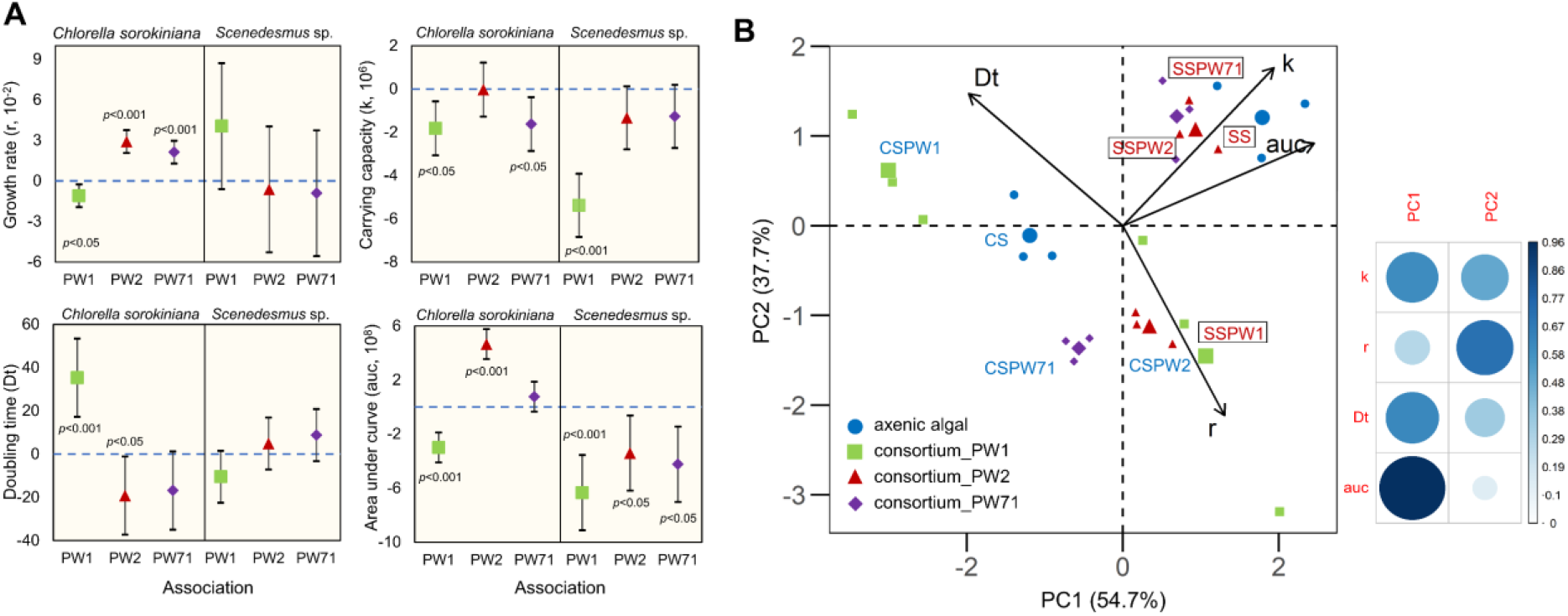
Assessment of algal growth parameters in algal-bacterial consortium under iron-limiting conditions. **A** Confidence interval plots of various algal growth parameters in consortium with respect to axenic growth (horizontal blue dashed line). Under iron stress, *Ralstonia pickettii* PW2 significantly increased growth rate (r) and area under curve (auc) of *C. sorokiniana* w.r.t. axenic culture. On the contrary, bacterium *Serratia plymuthica* PW1 showed significant reduction in growth rate, carrying capacity, and area under curve matrices of *Chlorella*. **B** PCA biplot of *C. sorokiniana* in consortium with PW1, PW2 and PW71 examining growth rate (r), doubling time (Dt) carrying capacity (k) and area under curve (auc). The biplot separates the *Chlorella sorokiniana* in consortium (consortium_PW2) from axenic algal setup on the basis of growth rate (r). Whereas, bacterium *Serratia plymuthica* PW1 increased the doubling time of the algae (consortium_PW1). The growth of algae *Scenedesmus* (SS), on the other hand, was not significantly different when co-cultured with all bacterial strains except for *Serratia plymuthica* PW1.

*S. plymuthica* PW1 exerted an antagonistic effect on *C. sorokiniana* as indicated by its significant increase in its doubling time (Dt) (*p*=0.009)with lower growth rate (r), carrying capacity (k), and area under curve (auc), comparing with the axenic culture of microalga (Fig. 3A). However, the lack of significant difference in growth parameters of *C. sorokiniana* co-cultured with *S. liquefaciens* PW71 indicates a neutral relationship between the two (Supplementary Table S2). *S. plymuthica* PW1, *S. liquefaciens* PW71 and *R. pickettii* PW2 showed an antagonistic effect on *Scenedesmus* sp., though the degree of effect varied (Fig. 3A). Co-culture of *S. plymuthica* PW1 with *Scenedesmus* sp., for example, caused a significantly reduced carrying capacity (*p*=0.000) and area under curve (*p*=0.003) of *Scenedesmus* sp. (Supplementary Table S2). Among these bacteria, *S. plymuthica* PW1 was antagonistic to both the microalgae.

Biplot based on principal component analyses (PCA) of microalga growth parameters in different experimental setups explained the mutualistic, antagonistic, and neutral effect of siderophore-producer bacteria on the growth of *C. sorokiniana* and *Scenedesmus* sp. (Fig. 3B). In PCA bioplot, *C. sorokiniana* (CS) grown axenically and in consortium with *R. pickettii* PW2 (CSPW2) formed separate groups based on the growth rate (r). Increasing growth rate (r) of microalgae with the consortium in iron-limiting condition confirms the benefits to microalgae due to iron made bioavailable by a siderophore-producing bacterium [55]. Bioavailable iron continues to accelerate microalgal growth until the iron exhausts at the stationary phase (Supplementary Fig. S2B). Such an iron-dependent mutualistic interaction between previously non-associated algae and bacteria has been shown between *Dunaliella bardawil* and *Halomonas* sp., where *Halomonas* sp. facilitates the iron in exchange for microalgal DOM [41]. In a marine environment, the vibrioferrin siderophore produced by *Marinobacter* sp. accelerates *Scrippsiella trochoidea* growth [19]. Similarly, *Idiomarina loihiensis* RS14 siderophore promotes the growth of freshwater alga *Chlorella variabilis* [56]. On the contrary, *C. sorokiniana* grown in co-culture with *S. plymuthica* PW1 (CSPW1) and as an axenic alga (CS) separated on the biplot due to distinct ‘doubling time (Dt)’, which is also supported by the slower growth in consortium due to antagonism by a bacterial partner as observed from algal growth curve (Fig. 2A).

*Scenedesmus* had a faster doubling time than *Chlorella*, therefore, grouped along with area under curve (auc) and carrying capacity (k) in the PCA biplot (Fig. 3B). In PCA biplot, *Scenedesmus* sp. from three setups, i.e., axenically grown (SS), co-cultured with *R. pickettii* PW2 (SSPW2), and *S. liquefaciens* PW71 (SSPW71) were grouped along with the growth variables, carrying capacity (k) and area under curve (auc) metric. As ‘auc’ metric explains 54.7% of variations in the grouping, it represents the dominant variable or Principal Component 1 (PC1) (Fig. 3B). Thus, the presence of bacterial strains *R. pickettii* PW2 and *S. liquefaciens* PW71 showed neutral interactions with *Scenedesmus* sp. as also observed from algal growth curves (Fig. 2A). *Scenedesmus*, however, was affected by co-culturing with *S. plymuthica* PW1 (SSPW1) in comparison to axenic growth (SS), as both were plotted separately due to reduced algal growth in the consortium (Figs. 2B, 3B). On the other hand, the growth rate (r) forms the PC2, which explains 37.7% of the variations in the grouping and explains the faster growth of *C. Sorokiniana* in consortium with siderophore-producing *R. pickettii* PW2 under iron-limiting conditions.

The HPAEC analyses of exopolysaccharides (EPS) of *C. sorokiniana* detected galactose (0.03±0.0 gL^-1^gcell^-1^; 52%) as a dominant monosaccharide besides glucose (0.01±0.0 gL^-1^ gcell^-1^, 20%), mannose (0.01±0.0 gL^-1^ gcell^-1^; 20%), arabinose (4%), and rhamnose (4%) (Fig. 2C and Supplementary Fig. S3A). *S. plymuthica* PW1 grew well in all five monosaccharides (0.1%), being galactose as the most preferred carbon source (Fig. 2D). *R. pickettii* PW2 showed a preference for galactose followed by glucose and mannose, though the growth promoted was relatively lower than that observed in *S. plymuthica* PW1. Sugar composition in microalgal EPS, governs microalgal-bacterial interaction [57]. In *C. sorokiniana*, galactose has been previously reported as a dominant monosaccharide (67%). Galactose also serves as a signaling molecule in bacteria, and its presence in EPS has been hypothesized to extend stationary phase in microalga *Botryococcus braunii* [58]. In contrast with *Chlorella, Scenedesmus* sp. showed a different sugar profile with glucose (37%), mannose (32%), and rhamnose (12%) as major monosaccharides (Fig. 2A). Bacteria grew well on *Chlorella* exudates rich in galactose than the exudates of *Scenedesmus* sp. rich in glucose and mannose (Supplementary Fig. S2B), suggesting the sugar profile of EPS govern the microalgal-bacterial association.

EPS-associated mono- and oligo-saccharides act as chemotactic molecules in microalgae to attract bacteria rather than serving as a source of carbon for bacterial growth [59]. Bacteria secreting exoenzymes hydrolyzing high molecular weight polysaccharides for continued carbon can sustain such associations. *R. pickettii* PW2 showed high growth in co-culturing with *C. sorokiniana*, suggesting its EPS preference (Fig. 2B). Though the commensal association between *C. sorokiniana* and bacteria with a high percentage identity to *R. pickettii* has been shown under nutrient-sufficient photoautotrophic conditions [60], our study suggests *Chlorella* EPS serves both as a source of DOM and chemotactic molecule, which governs the mutualistic association between them under the iron limitation. Furthermore, quorum sensing in microbes regulates the dynamic social behavior of bacteria and algae in the presence of ‘public goods’ like sugars and siderophore-chelated iron, a subject of the previous investigations [61]. Metagenomics also shows co-occurrence of *R. pickettii* with microalgae *Botryococcus braunii* [62], suggesting a wider environmental co-existence of the bacteria with microalga found in fresh and brackish waters.

Under experimental conditions, *S. plymuthica* PW1 showed ~10 times more growth than *R. pickettii* PW2 when co-cultured with *C. sorokiniana* (Fig. 2B). Antagonism between *S. plymuthica* PW1 and *C. sorokiniana* could be attributed to its aggressive growth (Fig. 2A). Also, *S. plymuthica* PW1 grew in all five sugars, suggesting a generalist and competitive life strategy (Fig. 2D). *Serratia* is also known to secrete serine protease, which lyses algal cells, providing a competitive advantage [63].

### Bioavailable iron influences dye degradation by *Chlorella sorokiniana*

To ascertain the significance of biologically available iron from bacterial siderophore, the dye degradation potential of *C. sorokiniana* was analyzed in axenic and consortium culture in iron-deficient and -sufficient conditions. EDTA-chelated iron increased dye decolorization potential of *C. sorokiniana* after 24 h (Fig. 4A). In treatment setups without EDTA-chelated iron, co-culture of microalga and bacteria significantly reduces half-life of AB1 dye compared to the axenic algal setup (*p*=0.000; consortium: half-life = 65.40±0.5 h; axenic algal: half-life = 120±9.4 h) (Supplementary Table S3 and Table S4). However, in the setups with EDTA-chelated iron, the half-life of AB1 lacks any significant difference between the experimental setup with the consortium and axenic algae (*p*=0.806; consortium: half-life = 15.27±2.5 h; axenic algal: half-life = 21.59±1.9 h) (Fig. 4A). In an experimental setup without EDTA-chelated iron, the dye degradation follows Simple First Order (SFO) kinetics [64]. In contrast, with EDTA-chelated iron, the degradation follows a First Order Multi-Compartment (FOMC) (Fig. 4A and Supplementary Fig. S4). FOMC represents a biphasic degradation indicating an initial steep decline in dye concentration followed by a relatively slower degradation. Co-culturing of siderophore-producer *R. pickettii* PW2 and *C. sorokiniana* significantly increased dye degradation by microalga only in the absence of EDTA-chelated iron (Fig. 4A). Bacteria produce siderophore in low iron conditions [65]; however, EDTA being a strong chelator, significantly increases available iron (Fig. 4A and Supplementary Table S4). Bacterial presence, therefore, lacks any significant effect on algal dye degradation due to the high bioavailability of iron (*p*=0.807) (Fig. 4A and Supplementary Table S4). Thus, *R. pickettii* PW2 increases the dye degradation potential of *C. sorokiniana* only under iron-limiting conditions. In contrast, axenic bacteria lack detectable dye degradation suggesting a significant contribution of a microalgal partner in dye degradation in the consortium.

**Fig. 4.**
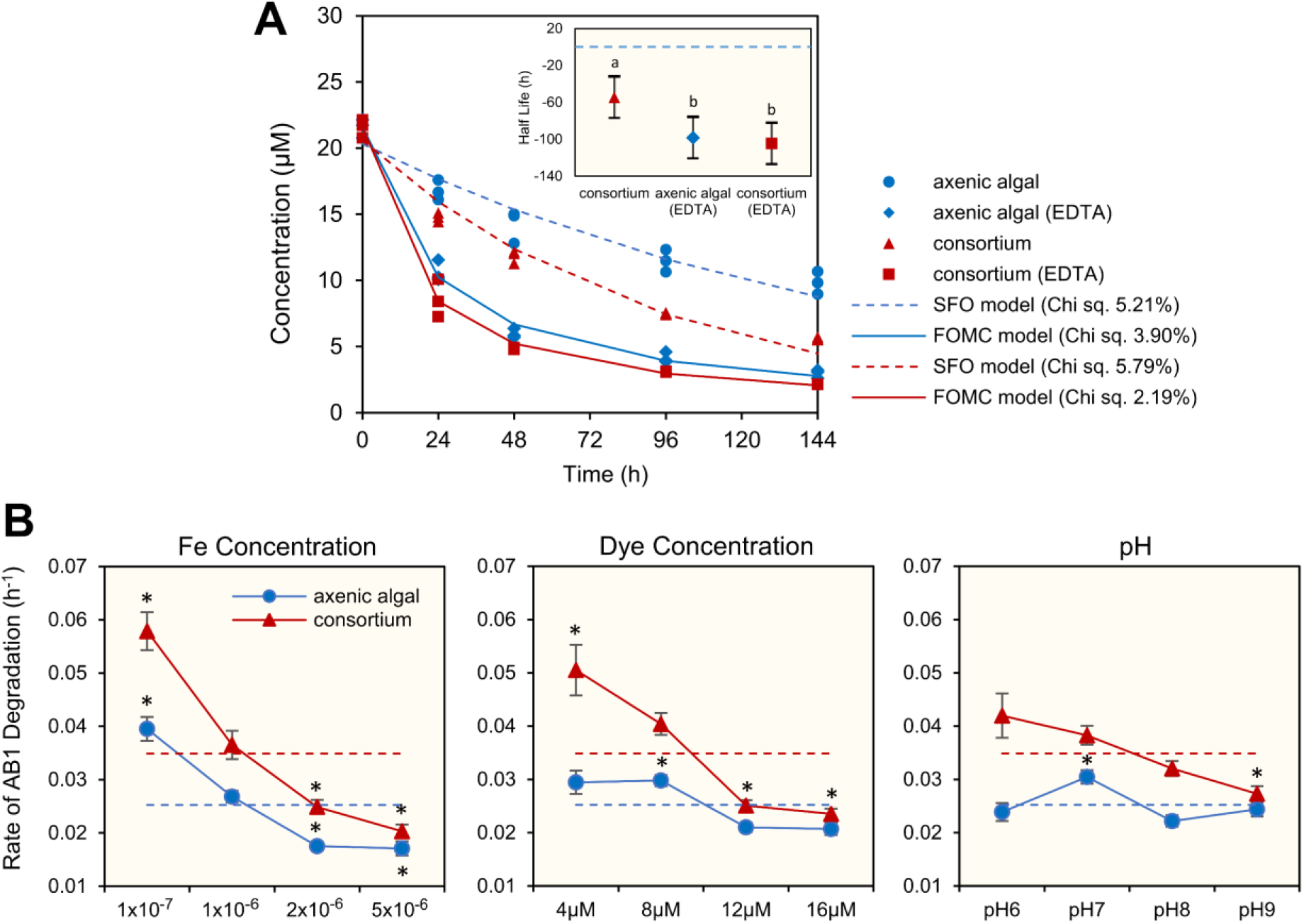
Dye degradation assessment of algal-bacterial consortium. **A** Degradation of Acid Black 1 (AB1) textile dye followed a Single First Order (SFO) kinetics without EDTA and biphasic First Order Multi-compartment (FOMC) with EDTA. Concentration of AB1 after incubation with *C. sorokiniana* in axenic and consortia set ups. Inset; blue horizontal dashed line represents half-life in axenic algal setup without EDTA supplemented iron. Comparison of half-life with respect to culture without EDTA supplementation is denoted by ‘a’ and ‘b’ grouping after Tukey’s post hoc test. **B** Analysis of Taguchi’s orthogonal array using a multiple linear regression model to investigate the impact of iron concentration, dye concentration and pH on AB1 degradation. Concentration of iron has the highest effect on the rate of AB1 degradation followed by dye concentrations. The ‘*’ symbol represents the significant difference of the means from the coefficient (dashed line; Supplementary Table S7).

To determine the role of biotic and abiotic factors in dye degradation, the factors relevant to industrial wastewater were chosen in designing 32 experiments with two setups, i.e., axenic microalgal culture and microalgal-bacterial consortium (Supplementary Fig. S5 and Table S5). Using L_16_ (4^3^) orthogonal array design, the dye degradation rate was calculated. L_16_ (4^3^) design was used to test varying Fe^3+^ and dye concentration and pH level. The Fe^3+^ concentration was kept lower or higher than 1×10^-6^ M, a concentration known to induce iron-starvation due to variation in the equilibrium between intra- and extracellular Fe concentration, thus necessitating bacterial siderophore production [33, 66]. pH determines the iron solubility and growth of microbes [67], whereas the concentration of AB1 dye (substrate) affects the enzymatic activity and the rate of dye degradation.

Multiple regression suggests the Fe^3+^ concentration (delta value; axenic algal = 0.02, consortium = 0.03) as a primary factor governing the rate of dye degradation followed by the azo dye concentration (delta value; axenic algal = 0.01, consortium = 0.02) (Fig. 4B, Supplementary Table S6 and Table S7). The delta value in Taguchi’s orthogonal design takes all the factors individually to determine the difference between the highest and lowest value of the average response variable. Therefore, a higher delta value of a particular factor represents a significant effect of variation in the level of the factor. Changing the concentration of Fe^3+^ led to a considerable variation in the rate of AB1 degradation in both axenic algal (*p*=0.001) and consortium (*p*=0.002) setups (Fig. 4B). In both the experimental setups, the rate of AB1 degradation was inversely proportional to the concentration of Fe (Fig. 4B). Consortium showed an enhanced average rate of AB1 degradation (0.04 h^-1^) as compared with axenic cultures (0.03 h^-1^) (Fig. 4B). Siderophore-producer bacteria increase dye degradation potential of microalga at 1×10^-7^ and 1×10^-6^ M Fe, whereas at a higher Fe^3+^ concentration, the bacterial effect remains neutral.

The dye concentration shows an inverse relationship with the rate of dye degradation in both axenic algal (*p*=0.028) and consortium (*p*=0.008) setups; however, it reached up to 60% even at high dye concentrations (Fig. 4B and Supplementary Table S5). At a high concentration, the dye molecules compete for electrons generated by the azoreductase-mediated enzymatic pathway at the microbial membrane [68], reducing dye degradation rate. Similarly, the siderophore-producer bacteria increased the rate of dye degradation at low dye concentrations only (Fig. 4B, Supplementary Table S6 and Table S7). Dye degradation in microbes is a non-growth associated extracellular process driven by plasma membrane-bound azoreductase, a non-specific oxidoreductase [48]. Azoreductase facilitates the transfer of electrons from microbial cells to electron-deficient azo bond (-N=N-), which reduces azo dyes into colorless aromatic amines via a two-cycle transfer of electrons following a ping-pong bi-bi mechanism [29, 68]. Azoreductase is engaged in oxidoreductase reaction with extracellular intermediates such as redox mediators like flavins which transfer electrons from within the microbial cell to extracellular azo bond. Therefore, the optimal ratio between the microalgal cells and dye molecules has a significant influence on catalytic efficiency and turnover number of enzymes for azo dye degradation. Contrary to this, the difference in the rate of AB1 degradation at varying pH was not significant (*p*>0.1) in both consortium and axenic algal setups (delta value; axenic algal = 0.01, consortium = 0.01) (Supplementary Table S6). Thus, as per the L_16_ orthogonal design, the variation in pH has a little effect on dye degradation potential by microalgae *C. sorokiniana* (Fig. 4B).

UPLC, FTIR, and LC-MS analysis confirm AB1 degradation by the microalgal-bacterial consortium (Fig. 5A). In chromatogram, the peak at retention time (RT) 2.617 corresponding to AB1 dye disappeared. The new peaks at RT 0.674, 0.944, and 1.944 appeared, which confirms the formation of less polar products on azo dye degradation. In the FTIR analysis, the disappearance of vibrational bands at 1,488 cm^-1^ and 1,282 cm^-1^ suggests azoreductase-mediated cleavage of azo bond (-N=N-) in AB1 dye (Fig. 5B) [1]. The disappearance of sharp intensive bands at 1,331 cm^-1^ and 642 cm^-1^ attributing to S=O and C-S bonds represents the removal of highly polar sulfo (R-SO_3_^-^) group in AB1 dye and formation of less polar byproducts. In the FTIR spectra of degradation products, the appearance of new vibrational bands at 1,652 and 1416 cm^-1^ (aromatic C=C), 1,589 cm^-1^ (-NH_2_), 3,452 cm^-1^ (-OH), and 1,579 cm^-1^ (-NO_2_), signifying the presence of aromatic hydrocarbons with nitro and amine substituted functional groups [1]. After that, the detection of degraded byproducts of AB1 via LC-MS confirms the azoreductase-mediated reductive cleavage of the dye (Supplementary Fig. S6 and Fig. S7). 4-Nitroaniline (RT 2.7 min, m/z 138.06) and naphthalene-1,2,8-triol (RT 2.45 min, m/z 176.1) are formed due to azoreductase-mediated symmetrical cleavage of the azo bond (Fig. 5C) [1]. Microbes further transformed 4-nitroaniline to 4-nitrophenol (RT 2.36 min, m/z 139.04), nitrobenzene (RT 2.33 min, m/z 122.04), and 4-aminophenol (RT 1.78 min, m/z 109.10). Naphthalene-1,2,8-triol was reduced to 1-naphthol (RT 1.7 min, m/z 144.9). The presence of catechol (RT 2.3 min, m/z 110.07), a central intermediate of aerobic biodegradation of benzene derivatives, suggests degradation via meta- or ortho-cleavage pathways [69].

**Fig. 5.**
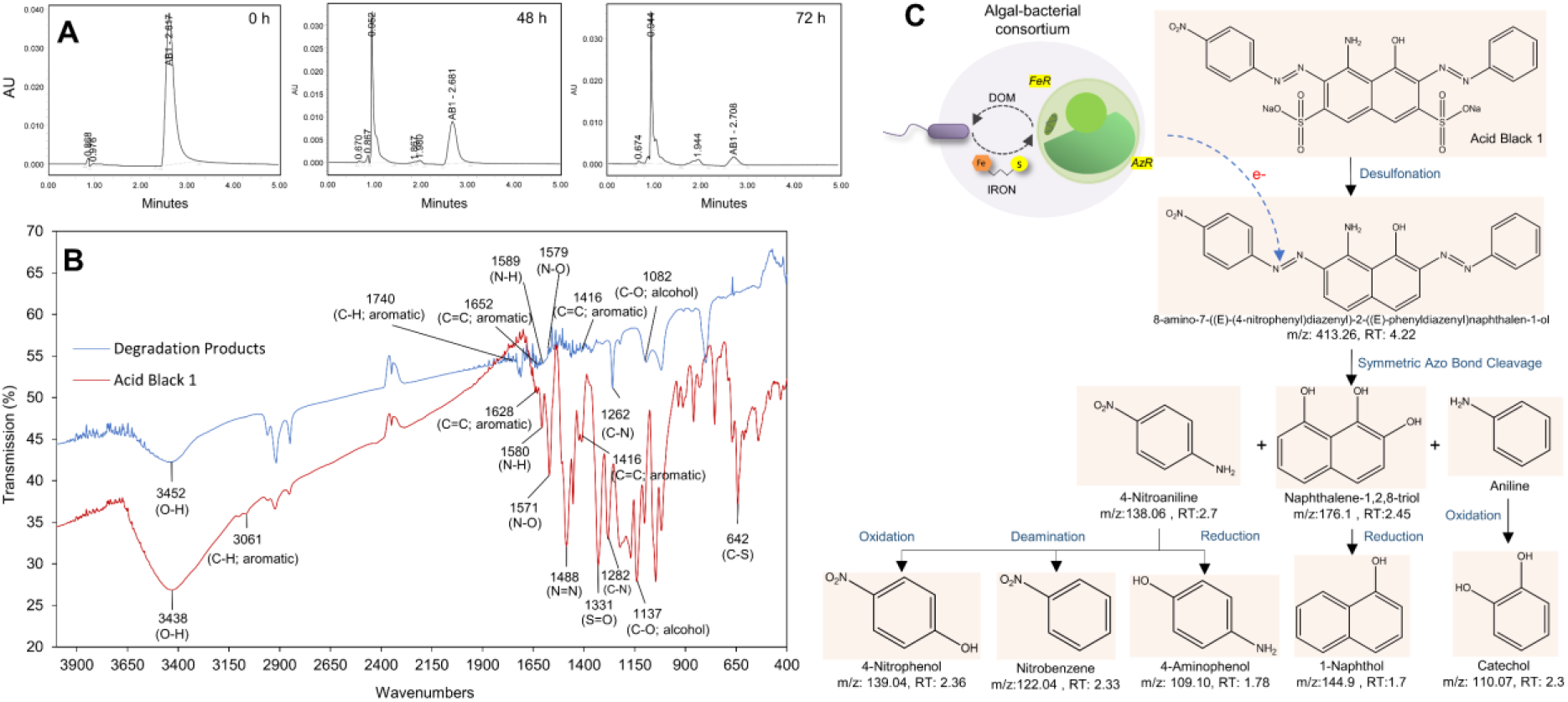
Analysis of AB1 degradation products. **A** UPLC analysis suggests the disappearance of the peak (RT 2.617) of Acid Black 1 (AB1) dye after treatment with algal-bacterial consortium. **B** The FT-IR spectra of Acid Black 1 dye and its degradation products after microbial treatment suggests the difference in the functional groups. **C** LCMS analysis of degradation products, and a suggested microbe-mediated biodegradation pathway of AB1 represents the azoreductase-mediated symmetrical cleavage of azo bond.

### Plasma membrane-bound ferrireductase enzyme influences algal dye degradation

We hypothesized that the increase in the rate of azo dye degradation with increased bioavailable iron has an association with higher activity of cellular azoreductase per cell or higher activity of azoreductase due to an increase in cell numbers. Therefore, axenic algal and algal-bacterial consortium were subjected to different Fe concentrations, and their effect on the algal cell growth, enzymatic activities, and iron uptake was assessed. With the increase in iron concentration, algal cell growth showed a considerable increase (Fig. 6A). In axenic cultures, *C. sorokiniana* growth varied with Fe concentration in the experimental setup, 2×10^-6^ M Fe: 8×10^7^ cells mL^-1^, 1×10^-6^ M Fe: 6×10^7^ cells mL^-1^; and 1×10^-7^ M Fe: 1.4 ×10^7^ cells mL^-1^. At lower Fe concentrations, *R. pickettii* PW2 enhances the growth of *C. sorokiniana*. The effect of the siderophore-producing bacteria in enhancing algal growth at lower Fe concentrations was also significant on growth parameters, such as growth rate (r): *p*=0.006 for 1×10^-7^ M Fe, *p*=0.028 for 1×10^-6^ M Fe; carrying capacity (k): *p*=0.025 for 1×10^-7^ M Fe; and *p*=0.017 for 1×10^-6^ M Fe, and area under curve (auc): *p*=0.003 for 1×10^-7^ M Fe; and *p*=0.001 for 1×10^-6^ M Fe (Supplementary Fig. S8, Table S8, and Table S9). However, at 2×10^-6^ M Fe, microalgae, grown axenically and in the consortium, lack a significant difference in their growth parameters (*p*=0.157 for r, *p*=0.551 for k, and *p*=0.444 for auc). Also, NO_3_-N concentration in axenic algal and algal-bacterial setups differ insignificantly even after 14 days, suggesting the cells are not nitrogen depleted.

**Fig. 6.**
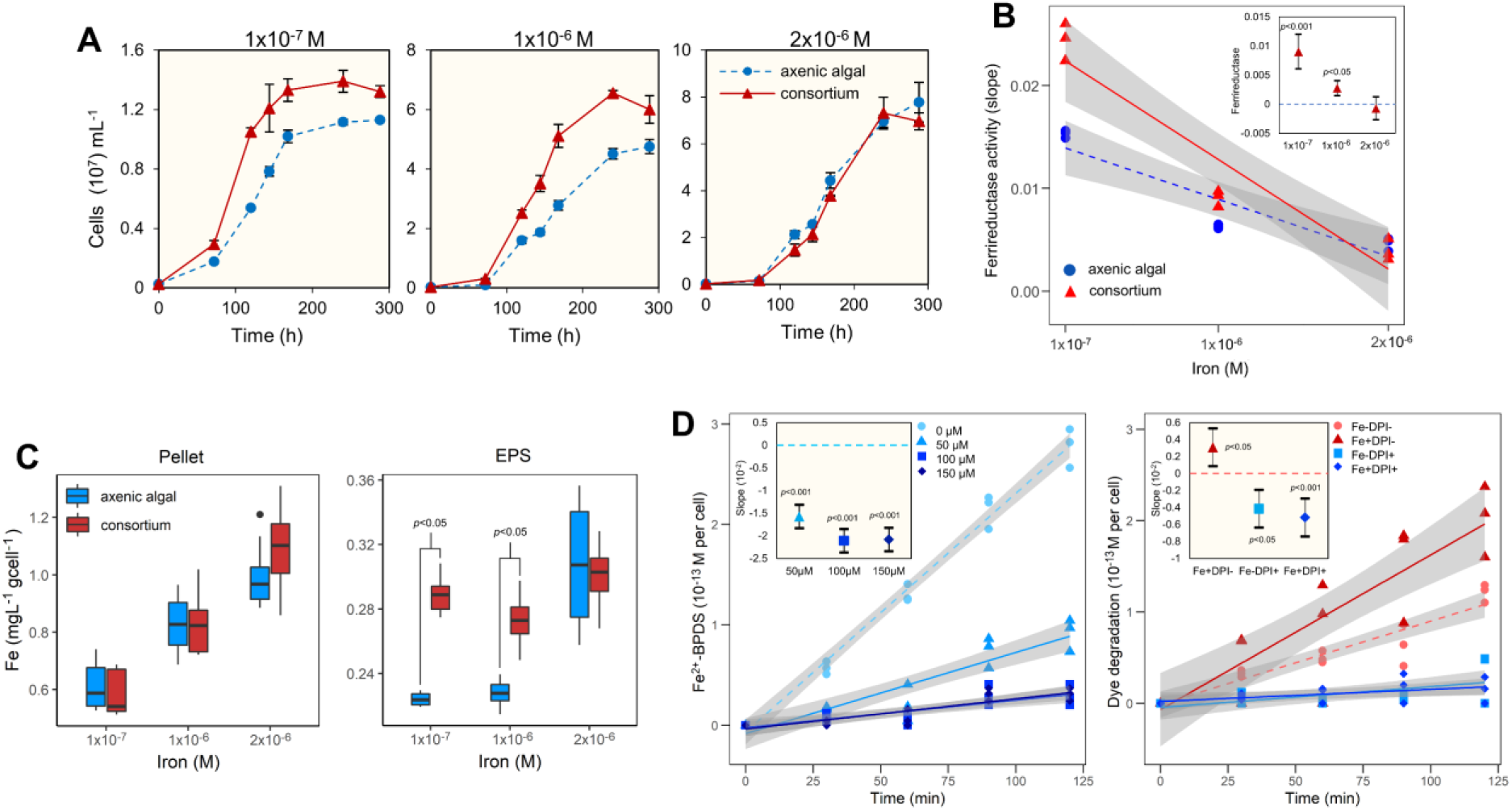
Impact of iron concentration on the growth and enzyme activity of *C. sorokiniana*. **A** Cell counts from algal-bacterial co-culture under three different iron concentrations. **B** Assay of extracellular ferrireductase activity of algae under different iron concentrations suggests bacteria-mediated significant increase in enzyme at lower Fe concentration (inset; confidence interval plot). **C** ICP-MS analysis of iron concentration in cell pellet and EPS under different iron concentrations. Algae accumulate more iron within EPS in presence of bacteria (*t*-test, *p*<0.05). **D** Ferrireductase assay conducted in the presence of Diphenyleiodonium (DPI) to assess extracellular iron-reductive pathway and azoreductase assay to assess azo dye degradation in *Chlorella sorokiniana*. Here, Fe-DPI-, Fe+DPI-, Fe-DPI+, and Fe+DPI+ represents different treatment setups and ‘+’ and ‘-’ denotes presence and absence. Pre-treatment of algal cells with 50 μM DPI significantly reduced extracellular ferrireductase activity and azoreductase activity.

Though algal cell density increases at high Fe concentration, the dye degradation rate significantly decreases (Figs. 4b, 7a). In *C. sorokiniana*, membrane-bound ferrireductase activity reduces with increasing iron concentration (Fig. 6B) [70]. Also, siderophore-producer *R. pickettii* PW2 significantly increased ferrireductase activity in microalgae at lower Fe concentrations (1×10^-7^ M Fe: *p*=0.001; and 1×10^-6^ M Fe: *p*=0.003) (Supplementary Table S10). Ferrireductases reduce Fe^3+^-siderophore complexes to release bioavailable Fe^2+^, which can be directly engulfed via an endocytosis-mediated non-reductive pathway [19, 71]. In contrast, despite a considerable increase in microalgal growth in the consortium at a high Fe concentration, the ferrireductase activity lacked any significant increase (Fig. 6B), which could be a potential shift in an iron-uptake mechanism. At higher Fe concentrations, bacteria use ferric iron via direct diffusion across the cell membrane than via high-affinity iron uptake by producing siderophore and chelating iron [33, 39]. Similar to bacteria, at high iron availability, microalgae follow a non-reductive direct iron-uptake pathway triggered by the difference in intracellular and extracellular iron concentration (Supplementary Fig. S9) [72, 73].

**Fig. 7.**
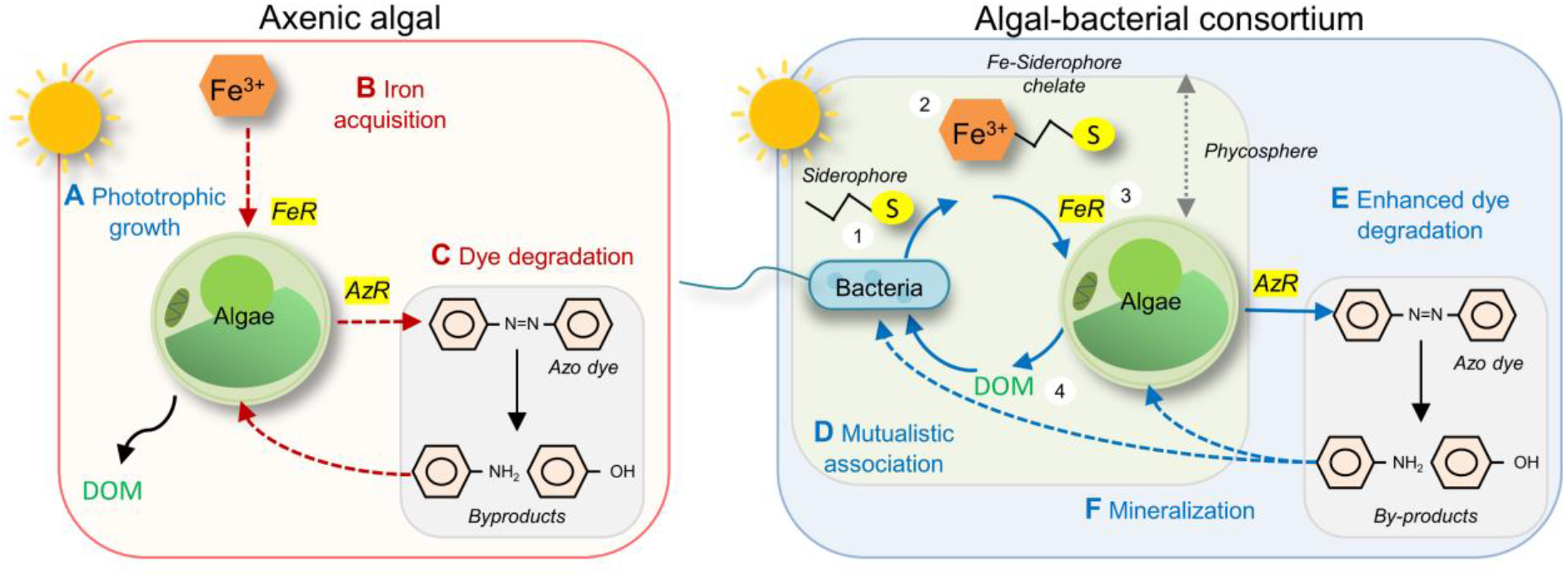
Proposed advantages of working with an algal-bacterial consortium in comparison to axenic algal treatment setups for remediation of dyes. **A** In a phototrophic textile wastewater treatment setup, **B** algae have to perform multiple tasks like iron acquisition, and **C** extracellular degradation of toxic dyes which leads to metabolic burden (*dashed red lines*) and affects algal growth. Algae has a poor iron uptake mechanism, however, a consortium between algae and siderophore producing bacteria, will increase the bioavailability of iron for algae and reduce the metabolic burden (*blue solid lines*’). **D** The mutualistic association between algae and bacteria takes place in the algal phycosphere (aquatic analogous of rhizosphere) which allows the exchange of micronutrients like iron, vitamins, Dissolved Organic Matter (DOM) [19]. The bacterial-secreted siderophores chelates non-bioavailable Fe^3+^ and increased the iron bioavailability for ferrireductase (FeR) mediated uptake (steps 1 to 3 in figure). Under iron limiting conditions, the siderophore-mediated increase in iron bioavailability also increases ferrireductase activity, and thereby, algal growth. Iron is an important macronutrient required for growth, enzymatic reactions, photosynthesis, and nitrate reduction in algae. Algae, on the other hand provides DOM for sustenance of bacteria (step 4) [21]. **E** From the study on *Chlorella sorokiniana* and *Ralstonia pickettii* PW2, the increase in bioavailability of iron also influenced the extracellular azoreductase (AzR) mechanism. The experimental evidence from this study suggests that the bacteria enhanced the algal ferrireductase and azoreductase activity, thus highlighting the potential link between these two enzymes [75]. **F** Therefore, a consortium can decolorize the dye and further mineralize the degradation products. The microbial consortium with autotrophic algae and heterotrophic bacteria will require minimum input for growth and will provide a sustainable bioremediation strategy.

Our results demonstrate an association between siderophore-mediated bioavailable iron and algal ferrireductase activity. In contrast with axenic culture, microalgal cell surface EPS accumulates iron significantly more when co-cultured with bacteria (*t*-test; *p* < 0.05) (Fig. 6C) but only at low Fe concentrations: 1×10^-7^ M and 1×10^-6^ M, indicating iron accumulation is a stress response (Supplementary Fig. S9) [72]. Algae do not produce siderophores; however, they increase iron in the phycosphere by biosorption and chelation onto extracellular polymeric substances, including mono- and polysaccharides [20]. Iron accumulation at the surface acts as a signal for ferric-assimilating proteins (FEA1 and FEA2) to assimilate the chelated iron for intracellular uptake via ferrireductase-dependent reductive pathway (Supplementary Fig. S9), which has also been shown in marine microalga *Chromera velia* [74], and in *Chlorella sorokiniana* UTEX 1602 (Supplementary Table S13 and Table S14).

Iron serves as an essential micronutrient in microbes; however, the association between bioavailable iron and microbial efficacy to degrade azo dye has not been explored previously. To examine the effect of ferrireductase, microalgal cells were pretreated with 50 μM Diphenyleneiodonium (DPI), a known inhibitor. DPI prevents the transfer of an electron from ferrireductase (flavohemoproteins; Fre1) to the Fe-chelate, thereby inhibiting Fe^3+^ to Fe^2+^ reduction, thus, inhibiting ferrireductase activity (*p*=0.00) (Fig. 6D and Supplementary Table S11) [49, 50, 73]. DPI (50 μM) significantly inhibited microalgal azoreductase both in the presence (Fe+DPI+) (*p*=0.008) and absence (Fe-DPI-) (*p*=0.035) of iron (Supplementary Table S12). However, *C. sorokiniana*, if not pretreated with DPI (Fe-DPI-), retained significant azoreductase activity, which was further increased after iron supplementation (Fe+DPI-) (*p*=0.012) (Fig. 6D). We report NAD(P)H-dependent azoreductase in *C. sorokiniana* lacking sensitivity to oxygen in complete photoautotrophic culture conditions [29], which adds to its benefits for industrial sustainability.

In *Saccharomyces cerevisiae*, an association between azoreductase and ferrireductase activities has been shown [75]; however, such association has never been explored in microalgae. Azoreductase in *S. cerevisiae* shows a high dependency on the Fre1p component of metalloregulator ferrireductase (FRE1). Transmembrane ferric-chelate reductases having high sequence similarity have also been reported in green microalga *C. sorokiniana* UTEX 1602 and *Chlamydomonas reinhardtii* (Supplementary Fig. S10, Table S13, and Table S14). Taken together, we propose that the ferrireductase-mediated reductive pathway is vital for azoreductase in *C. sorokiniana*, though it requires further investigation in understanding the exact role of extracellular iron-concentration in driving the oxidoreductases enzymatic machinery in *Chlorella*. Typically, textile wastewater bioremediation studies treat systems as black boxes. A poor understanding of the extracellular environment, especially the role of micronutrients, has posed a major limitation in their industrial translation. In this work, extracellular iron concentration not only influences the plasma-membrane bound ferrireductase pathway in *Chlorella*, which sustains such inter-kingdom mutualism but also azoreductase-mediated dye degradation.

Recent photobioreactor-based comprehensive studies on algal-bacterial respiration, COD removal, and nitrification have further highlighted the potential of such inter-kingdom symbionts in replacing the non-specific bacterial processes in industries [30]. Bacterial azoreductases, which in general are oxygen-sensitive, require a complex two-stage design to integrate into the current industrial infrastructure. In contrast, *Chlorella* has been reported to modify extracellular oxygen in contrasting dark (oxygen-deprived conditions favoring dye degradation) and light (oxygenated conditions favoring aromatic amines degradation) cycles accelerate dye degradation [12], and provide oxygen for bacterial growth and COD removal in municipal wastewater [30]. In our work, the *Chlorella-Ralstonia* consortium showed a higher algal growth rate (~ 2 times) and dye degradation rate (~ 2 times) compared to monocultured alga. The consortium also overcame iron limitation and carbon availability and performed under varied pH (pH 6 to 9) and dye concentrations. Therefore, we propose that such algal-bacterial mutualistic associations could form a stable and robust bioremediation system that overcomes the current limitations of biological designs for textile industry wastewater treatment.

In conclusion, a consortium of *Chlorella sorokiniana* and *Ralstonia pickettii* PW2 represents a symbiotic association based on the exchange of specific limiting nutrients (Fig. 7). *C. sorokiniana* receives iron from *R. pickettii* PW2 in exchange for dissolved organic matter to develop a barter system. Under iron stress, bacterial siderophore ensures iron availability promoting algal growth rate and extracellular dye degradation. Therefore, a bacterial-algal association has the potential for wastewater treatment under iron-limiting conditions compliant with green chemistry principles. We report the transmembrane ferrireductase activity in *C. sorokiniana*, which plays a crucial role in a reductive iron-uptake pathway involving azoreductase. Bioavailable iron regulates the co-expression of the two oxidoreductase pathways. Microalgal-bacterial consortium works under photoautotrophic conditions would provide a self-sustainable alternative to current microbial dye remediation processes. Further studies on microalgal-bacterial mutualisms will help develop methods to overcome the limitations of micronutrient availability in industrial bioprocesses and bioremediation.

## Materials and Methods

### Culture media preparation, sample collection, and bacterial identification

The three freshwater algal strains were screened in this study, *Chlorella sorokiniana* (CCAP 211/8K), *Scenedesmus* sp., and *Oscillatoria animalis* (Sciento Ltd), were cultivated in Bold’s Basal Medium (3N-BBM+V; henceforth referred to as BBM) and maintained with EDTA chelated iron at 28°C under continuous white light (Fig. 1B). Two strains of the marine microalga *Phaeodactylum tricornutum* 1052/6 and 1055/1 were grown on sterile *f/2* medium [36]. FeCl_3_.6H_2_O was used as an iron source; however, 10-folds Na-EDTA was used as a chelating agent only when specified. The iron-abundant BBM is referred to as BBM+Fe (EDTA-chelated) and iron-deficient as BBM-Fe (not EDTA-chelated).

Textile wastewater was collected from the industrial area in Panipat, India (29.363121 N, 76.992971 E). The non-selective 10% tryptone soya agar (TSA) plates supplemented with 0.014% triphenyl tetrazolium chloride (TTC) was used to isolate bacteria from textile wastewater [37] (Fig. 1A). The morphologically distinct colonies were purified and identified as *Serratia plymuthica* PW1, *Ralstonia pickettii* PW2, *Stenotrophomonas rhizophila* PW3, *S. maltophilia* PW5, *S. maltophilia* PW6, *Serratia liquefaciens* PW71, and *Stenotrophomona rhizophila* PW72 using 16S rRNA sequence analyses (Table 1) [38]. Genomic DNA of bacteria was isolated and purified using Wizard Genomic DNA Purification Kit (Promega, USA). 16S rRNA gene was amplified using PCR with universal primers 27F (5’-AGAGTTTGATCCTGGCTCAG-3’) and 1492R (5’-TACGGYTACCTTGTTACGACTT-3’) for 30 cycles (94°C for 1 min, 50 °C for 1 min, and 72 °C for 2 min). The amplified products were analyzed onto 1% agarose gel (low EEO) and extracted and purified from the gel using the QIAprep Miniprep Kit (Qiagen, Netherlands). The 16S rRNA sequencing reactions were performed using 785F (5’-GGATTAGATACCCTGGTA-3’) and 907R (5’-CCGTCAATTCCTTTRAGTTT-3’). The homologous sequence was searched in the NCBI GenBank database using BLAST and DNA sequences of identified bacteria were submitted with accession numbers MW857264, MW857265, MW857266, MW857267, MW857268, MW857269, and MW857270. The isolates were revived in 10% TSB for further experiments (Fig. 1A).

### Bacterial siderophore production

Bacterial siderophore production was estimated using standard Chrome Azurol S (CAS) assays in modified Minimal Medium 9 (MM9) supplemented with casamino acid deferrated with 8-hydroxyquinoline, glucose, MgCl_2_, and CaCl_2_ [39, 40] (Fig. 1C). Siderophore production was ascertained as yellow/orange zone around bacterial colonies on CAS agar plate. Bacterial siderophore was quantified using standard CAS liquid assay using Desferrioxamine mesylate used as standard, and further categorized using Csaky’s assay (for hydroxamate type) with Desferrioxamine mesylate and Arnow’s assays (for catecholate) with 2’3-Dihydroxybenzoic acid as standards [39].

### Algal dye decolorization assessment

Microalgal potential to degrade Acid Black 1 dye (AB1, Sigma Aldrich; CAS: 1064-48-8) was assessed in their respective growth media (Fig. 1D). Ten-day old algal species cultures were preincubated in EDTA-chelated growth media for 48 h before adding filter-sterilized (0.2μ) 10 μM AB1 dye. Microalgal cultures were incubated at 28°C under continuous light for 72 h, and cell biomass was removed by centrifuging the culture at 5000×*g* for 5 min. The supernatant was used to estimate the dye decolorization by microalgal cultures by calculating the per cent dye removal as follows: [(initial OD_618nm_ - final OD_618nm_)/initial OD_618nm_]×100 [1].

### Algal-bacterial co-culturability assessment under iron limiting conditions

To analyze the symbiotic interactions in iron-limiting environment, dye-decolorizer microalgae (*Chlorella sorokiniana*/*Scenedesmus* sp.) were co-cultured with siderophore-producing bacteria (*Serratia plymuthica* PW1/*Ralstonia pickettii* PW2/*Stenotrophomonas maltophilia* PW5/*S. maltophilia* PW6/*S. liquefaciens* PW71), and their growth characteristics were determined. An overnight bacterial culture raised in 10% TSB was washed three times with sterile BBM-Fe. Bacterial suspension in BBM-Fe (OD_600nm_=0.3) was inoculated in microalgal exudates obtained by filter sterilization (0.2μ) of 7-day old algal culture raised BBM+Fe. The bacterial growth (OD_600nm_) was taken as a measure of the potential of bacterial isolate to use algal-derived dissolved organic matter (DOM) as a carbon source (Fig. 1E).

Bacterial isolates showing survival and growth in microalgal exudates were selected to ascertain their co-culturability [19] (Fig. 1F). Overnight cultures of bacterial isolates raised in BBM+Fe supplemented with 0.1% glucose were washed with sterile BBM-Fe, and 1 ml of bacterial suspensions (OD_600_ = 0.5) were inoculated in 20 ml of 1-day old cultures of microalgae raised in iron-deficient (1×10^-6^ M FeCl_3_) BBM-Fe media [33, 41]. Growth and morphological characteristics of both microalgal and bacterial co-inoculants were observed to determine the mutualistic, antagonistic, or neutral interactions. Bacterial and algal cells were determined periodically over 12 days on agar plates (CFUs) and under the microscope using a hemocytometer, respectively [19]. The algal growth curve was fitted in the logistic equation to determine the growth characteristics (growth rate ‘r’, carrying capacity ‘k’, doubling time ‘Dt’, and area under curve ‘auc’) using the growth prediction modeling package ‘*growthcurver*’ in R [42].

The significance of co-inoculation of siderophore producer bacteria on microalgal growth was statistically analyzed using One-Way ANOVA, Tukey’s post-hoc test, and Principal Component Analysis (PCA). Bacterial and microalgal isolates showing mutualistic associations were designated as a ‘consortium’ for further experiments on azo dye degradation (Fig. 1G).

### Characterization of carbohydrates in algal exopolysaccharides (EPS)

The exopolysaccharides (EPS) from *C. sorokiniana* and *Scenedesmus* sp. were fractionated using Dowex Marathon C cation exchange resin (Sigma Aldrich, USA) [43]. The algal cells from a 7-day old culture in BBM-Fe were washed and centrifuged at 2000×*g* for 3 min at RT. The cell pellets were resuspended in PBS buffer containing Dowex Marathon C (25g g^-1^) and gently mixed in a rotatory mixer (50 RPM) at 4°C for 1 h. Subsequently, the algal cells were centrifuged at 4000×g for 4 min, and the supernatant was subjected to overnight precipitation of exopolysaccharides in 70% ethanol (3:1 to supernatant) at 4°C [44]. Thereafter, the supernatant was centrifuged at 10,000×g for 10 min, and the pellet was collected for analyses by high-performance anion-exchange chromatography (HPAEC). The pellets were acid hydrolyzed with 72% H_2_SO_4_ at 121°C for 30 min in an autoclave. Monosaccharides present in hydrolyzed EPS were identified using Ion Chromatography System (Thermo Scientific Dionex ICS 5000+, UK) with AminoPac PA10 column (250 mm × 4 mm) and a pulsed amperometric detector. KOH (1 mM) was used for isocratic elution and separation at 0.25 mL min^-1^ for 25 min. Arabinose, fructose, galactose, glucose, mannose, and rhamnose were used as standards.

### Challenging algal-bacterial consortium under varying environmental conditions

To evaluate the performance of the microalgal-bacterial consortium in degrading dye, three experimental setups in BBM+Fe and BBM-Fe were conducted: (i) setup 1: axenic microalgal, (ii) setup 2: microalgal-bacterial consortium, and (iii) setup 3: axenic bacterial (supplemented with 0.01% glucose) (Supplementary Table S3). The microalgal cells, previously grown in BBM+Fe under static conditions at 28°C in continuous light, were washed and used in all the experiments. FeCl_3_ (1×10^-6^ M) was supplied to maintain the iron-deficient environment, whereas AB1 dye (20 μM) was filter-sterilized (0.2μ) in the experimental setups. The dye decolorization in different experimental setups was assessed for 144 h by using the linear regression equation obtained from the standard curve of AB1 dye [1]. The rate of AB1 degradation was calculated by fitting the concentration data in kinetic models using ‘*mkin*’ package in R [45].

To identify the factors governing the rate of dye degradation, a multi-factor study using Taguchi’s L_16_ (4^3^) orthogonal array design was developed. Thirty-two experiments were conducted in setups 1 (axenic algal) and setup 2 (algal-bacterial consortium) with varying concentrations of Fe, AB1 dye, and pH computed using Minitab ver. 19, as outlined elsewhere [46]. Details of variables/levels were as follows: Fe: 1×10^-7^, 1×10^-6^, 2×10^-6^, and 5×10^-6^ M; pH: 6.0, 7.0, 8.0, and 9.0, and (iii) dye: 4 μM, 8 μM, 12 μM, and 16 μM (Supplementary Table S5).

Microalgal cells were previously starved for iron in BBM-Fe media for 24 h at various pH levels as per the L_16_ design (Supplementary Table S5). For experiments in setup 2, a microbial consortium was developed by mixing an overnight culture of siderophore-producer bacterial isolate (OD=0.3) with microalgal cells (1 % v/v). The culture was spiked with different concentrations of iron (FeCl_3_.6H_2_O) and AB1 dye as per orthogonal array design and maintained under static conditions at 28°C with continuous light (Supplementary Table S5). Uninoculated BBM-Fe was kept as controls. Microbial cells were carefully separated, and culture media was sampled periodically over 48 h; dye concentration was determined in different experimental setups [1]. Also, the rate of AB1 degradation was calculated by fitting the concentration data in the first-order kinetic model using ‘*mkin*’ package [45, 47]. Variations in dye degradation in different setups were analyzed by multiple linear regression model using Minitab ver. 19, and the effect of varying levels of different factors on the rate of AB1 degradation was determined. The half-life of AB1 dye, the time at which 50% of dye gets degraded, and the chi-square (χ^2^) error level representing the goodness-of-fit of the kinetic model was also determined. A χ^2^ of <15 is considered as an acceptable fit [45].

### Assessment of algal ferrireductase, azoreductase activity, and iron concentration

Ferrireductase activity was determined in microalgae raised axenically and in consortium with siderophore-producer bacteria under different iron concentrations (FeCl_3_.6H_2_O: 1×10^-7^ M, 1×10^-6^ M, and 2×10^-6^ M). The growth characteristics (carrying capacity, growth rate, doubling time, and area under curve) of microalgal cells were observed using the ‘*growthcurver*’ package as described in section 2.4. After ten days of incubation, the microalgal cells were harvested, washed, and resuspended in PBS. An equal number of microalgal cells were maintained in all the assays [19]. The reaction mixture contained PBS with 130μM HEPES (pH 8.1), 10μM Fe (with 10-folds EDTA), and 100μM bathophenanthrolinedisulfonic acid or BPDS, i.e., Fe^2+^ chelator. The ferrireductase activity was estimated as the formation of the Fe^2+^-BPDS complex at OD_535nm_ using the extinction coefficient 22140 M^-1^cm^-1^.

Simultaneously, the cellular concentration of iron in microalgae was estimated using ICP-MS. The culture was centrifuged at 4000×*g* for 4 min, and the cells were removed, washed, and resuspended in sterile deferrated BBM. Microalgal cells were acid digested in 70% HNO_3_ and appropriately diluted for estimating iron content by ICP-MS (Bruker M90 ICP-MS). To estimate iron on the cell surface, microalgal EPS was extracted as described in section 2.5. The EPS was also digested in 70% HNO_3_ and diluted for estimating the Fe by ICP-MS. FeCl_3_ was used as a standard.

The azoreductase activity of microalgal cells was determined after 10 days of incubation, as described previously [48]. To confirm the expression of azoreductase in the microalgae during dye degradation, the microalgal cells from different setups were pretreated with ferrireductase inhibitor, i.e., Diphenyleiodonium (DPI) [49, 50]. The optimum concentration of DPI was determined by ferrireductase assay using microalgal cells pretreated with different concentrations of DPI (50, 100, and 150 μM) for 1 h. Excessive DPI was removed by a thorough washing of the cells, and azoreductase assay was carried out in a reaction buffer containing 50 mM phosphate buffer (pH 7.2) and 10 μM AB1 dye. The reaction was initiated by adding 0.5 mM NAD(P)H. The enzyme activity was determined by a decrease in OD_618nm_ over 240 min. Azoreductase activity was monitored at varying conditions of Fe (EDTA-chelated) and DPI concentration, i.e., Fe-DPI-, Fe+DPI-, Fe-DPI+, and Fe+DPI+. Linear regression model and One-Way ANOVA were used to analyze variations in the enzyme activities in different experimental setups.

### Assessment of microbe-mediated AB1 degradation pathway

The AB1 degradation pathway in an experimental setup with the microalgal-bacterial consortium was analyzed using analytical techniques. Microbial consortia cultivated in BBM-Fe were challenged with AB1 dye in the presence of 1×10^-6^ M FeCl_3_. Aliquots of culture were sampled aseptically at a regular time interval of the treatment by removing the microbial cells by centrifuging the culture at 10000×*g* for 15 min. The supernatant was filtered through a 0.2 μm filter, and the filtrate was analyzed by UPLC (Waters Acquity UPLC system, United States) using BEH C18 100mm× 2.1mm column fitted with a photodiode array detector. Acetonitrile with 0.2% formic acid in water (85:15 v/v) was used as a mobile phase, and the peaks were recorded at OD_595nm_. AB1 dye was used as a standard.

To decipher the possible dye-degradation pathway, AB1 degradation products in the different experimental setups were extracted using sequential solvent extraction and characterized and identified using different analytical techniques [1]. After dye decolorization, the microbial cells were pelleted (10,000×*g* for 15min at RT), and the supernatant was filtered through a 0.2 μ filter. The degraded products of the dye in the filtrate were extracted with an equal volume of ethyl acetate (Sigma Aldrich; HPLC grade) and dried over sodium sulphate (Na_2_SO_4_). Ethyl acetate was then evaporated using a rotary evaporator (Heidolph Laborota 4000, Germany) at 47°C under reduced pressure [1]. The dried metabolites were solubilized in a minimum volume of acetone subjected to FTIR analysis. Spectroscopically pure KBr was mixed with the samples (dissolved in HPLC grade acetone) in a ratio of 5:95 and analyzed at the mid-IR region (400-4,000 cm^-1^) of the spectrometer (Perkin Elmer Spectrum 100, United States). The biodegraded products of AB1 were identified using LC-MS analyses (Dionex Ultimate 3000) with acetonitrile: water (70:30) as a solvent system. The sample dissolved in HPLC grade methanol was injected in UHPLC (Hypersil Gold) using 5 μm, 100 cm × 2.1μm column for a run time of 0 to 20 min at a positive polarity (+1) mode. The mass spectra were recorded within a range of 100-1200 m/z with a maximum ion transfer time of 100 ms. Degraded products of AB1 dye were identified by analyzing mass fragment peaks and referring to NIST (National Institute of Standards and Technology), ChemSpider, and m/z cloud libraries and the existing literature.

## Acknowledgements

V.M. express gratitude towards the Department of Science and Technology, Technology Mission Division (Energy, Water & Others), India, for the financial support ‘‘Development of a novel single-stage…. environmental safety’’ under Optimal Water Use in Industrial Sector. V.M. and R.S.S also thank the Institute of Eminence for Faculty Research Project Grant. B.P. acknowledges support from the Winston Churchill Memorial Trust and the EPSRC Global Challenges Research Fund. D.R. thanks University Grant Commission, Government of India for providing Junior Research Fellowship, British Council, India, Newton Fund, UK, and Department of Biotechnology, Government of India, for Newton-Bhabha PhD placement award 2017-18. U.S. and P.P acknowledge the University Grant Commission for Junior Research Fellowship.

## Author contributions

D.R., V.M., B.P. and R.S.S conceived the experiments; D.R. performed the experiments; U.S., P.P., and A.F. contributed in siderophore method, multi-factor study, and ICP-MS study respectively; D.R. undertook data analysis; D.R., V.M., R.S.S. and B.P. wrote the manuscript.

## Data availability

The bacterial 16S rRNA sequencing data is available at the National Center for Biotechnology Information (NCBI) under submission accession number SUB9427182.

