## Supplementary Information for "Iron-dependent mutualism between *Chlorella sorokiniana* and *Ralstonia pickettii* forms the basis for a sustainable bioremediation system"

### Supplementary Figures

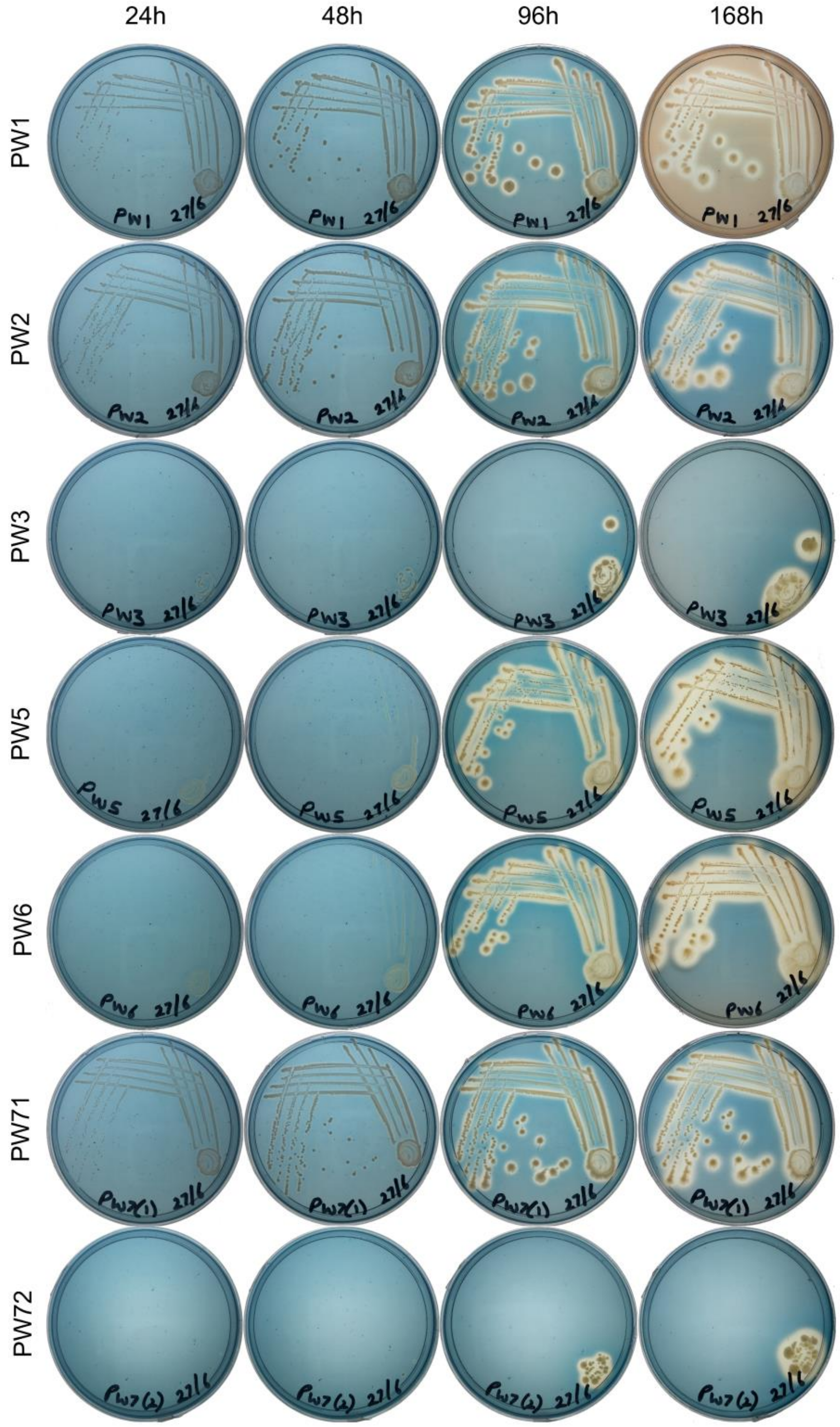


**Supplementary Fig. S1 The results from the CAS agar plate assay suggest the siderophore production in all 7 bacterial isolates though with varying growth in the iron limiting MM9 media.** The strains *Serratia plymuthica* PW1, *Ralstonia pickettii* PW2, and *Serratia liquefaciens* PW71 showed higher growth as the bacterial colonies appeared after 24 h of streaking. The bacterial isolates *Stenotrophomonas maltophilia* PW5 and *Stenotrophomonas maltophilia* PW6, though did not show immediate growth in the deferrated MM9 media, produced siderophore after 96 h of streaking. The bacterial strains *Stenotrophomonas rhizophila* PW3 and *Stenotrophomonas rhizophila* PW72 showed limited growth in MM9 media.

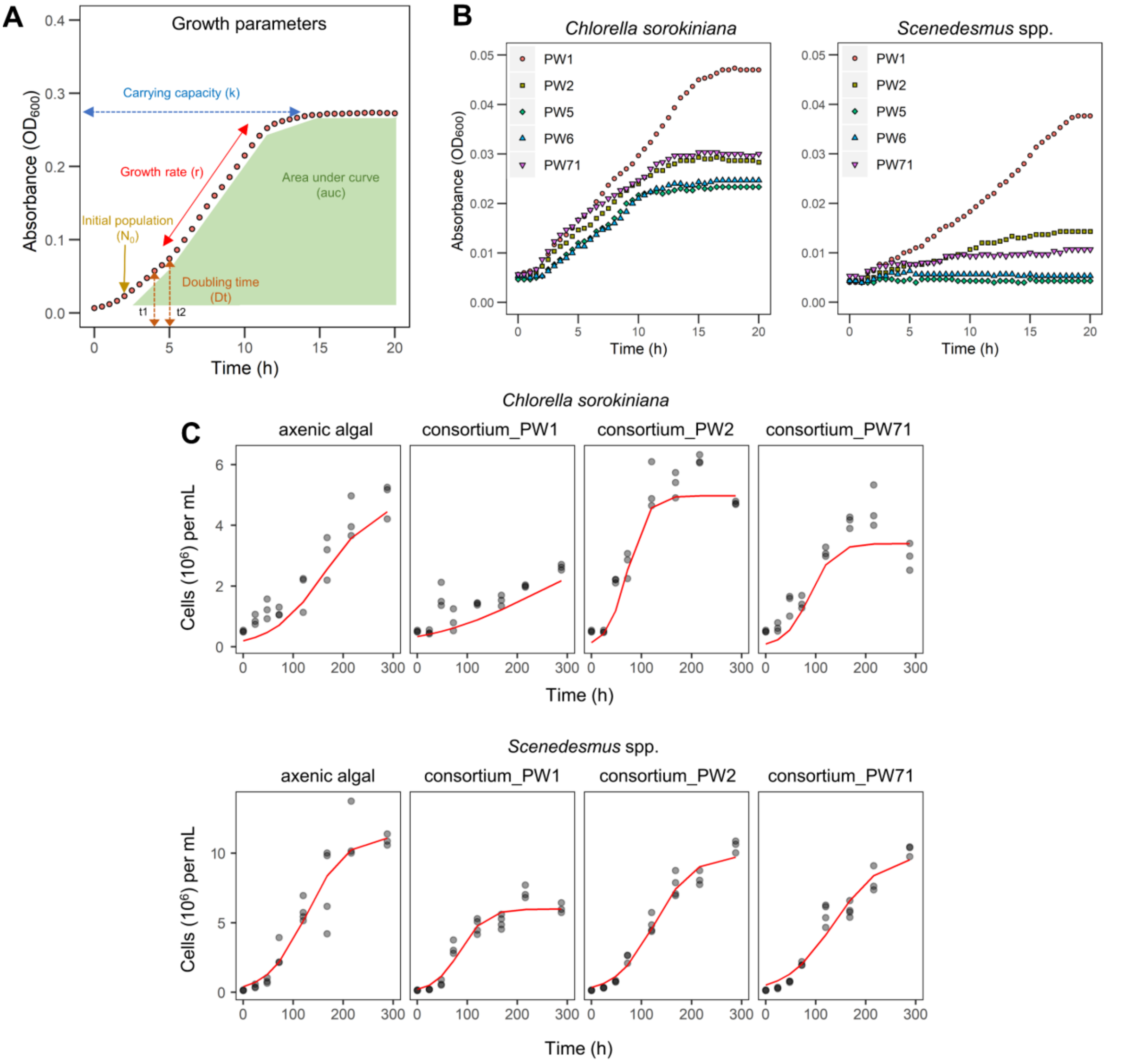


**Supplementary Fig. S2 Different growth parameters obtained from fitting the growth data using package '*growthcurver*' in R. A** The logistic equation Nt=k/(1+(k−N0/N0)e−rt) are explained as follows: the growth rate (r) represents an increase in the population of bacterial strains (per unit time) during the exponential growth phase. The doubling time (Dt) represents the time required for the microbial population to double in size (cell count) during the log phase. The carrying capacity (k) represents the maximum population size in the system, and area under curve (auc) is an integrative metric computed using the carrying capacity (k), growth rate (r), and initial population size ($N_0$) which represents the total area under the trapezoid-shaped growth curve. **B** Bacterial growth curves performed on exudates of *C. sorokiniana* and *Scenedesmus* sp. showed bacteria prefer exudates of *C. sorokiniana*. **C** The prediction model (red line) computed by fitting the observed algal growth data (grey dots) in the logistic equation using the prediction modelling package '*growthcurver*' in R [1].

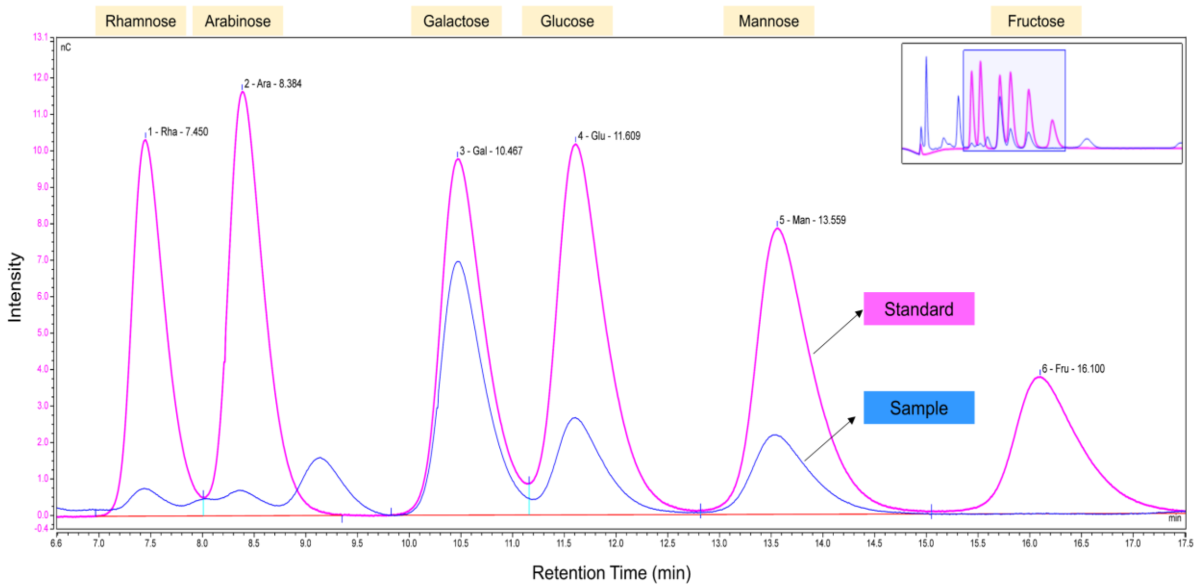


**Supplementary Fig. S3 Results of the anion-exchange chromatography and growth of bacterial strains in different carbon sources.** The chromatogram after high-performance anion-exchange chromatography represents the identification rhamnose, arabinose, galactose, glucose, and mannose in the EPS extracted from *Chlorella sorokiniana*.

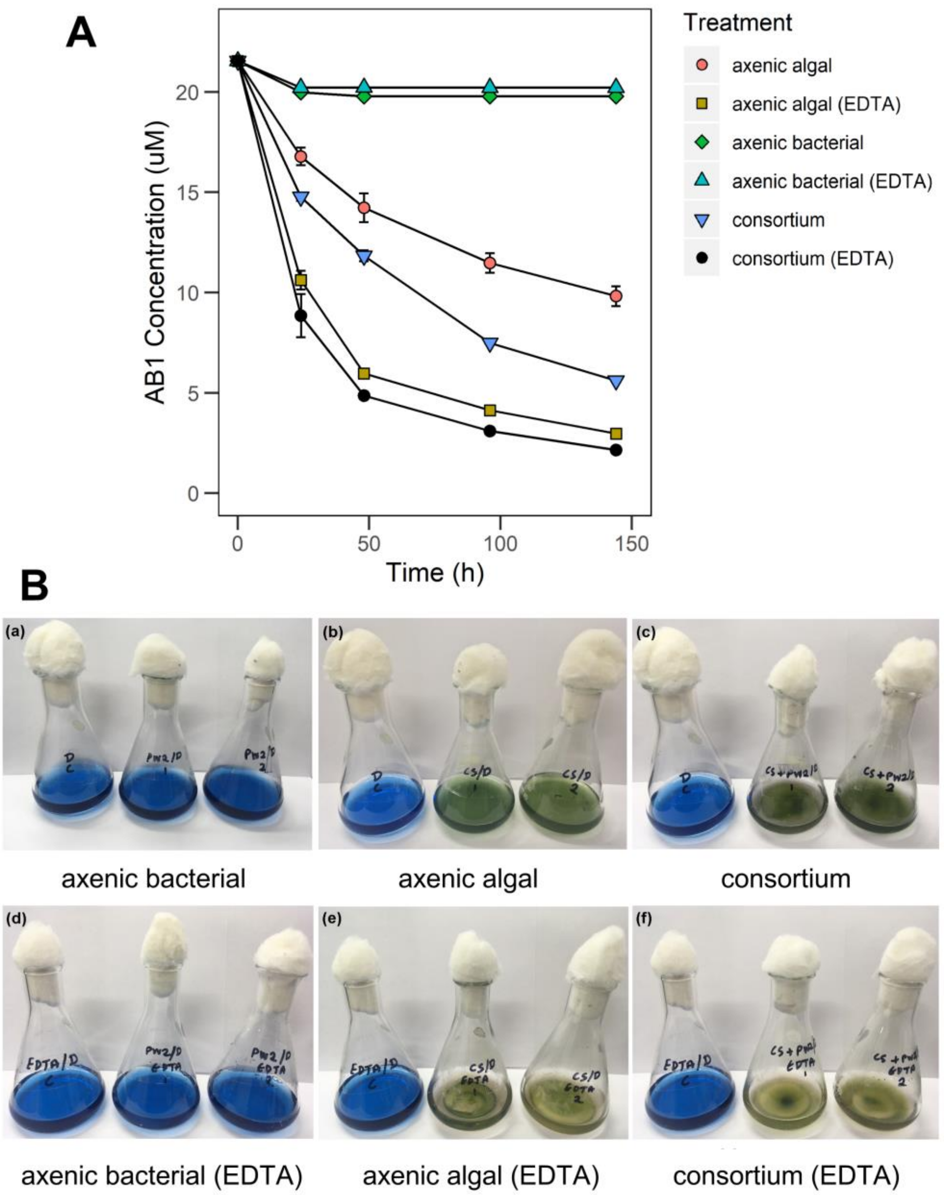


**Supplementary Fig. S4 The Acid Black 1 dye degradation under varied condition of iron bioavailability suggests a higher dye degradation in treatment setups with EDTA supplemented iron. A** The bacterial strain *Ralstonia pickettii* PW2 enhanced the algal dye degradation under culture conditions where iron was supplemented without EDTA (consortium). **B** The axenic bacterial setup did not decolorize dye even after 144 h of incubation.

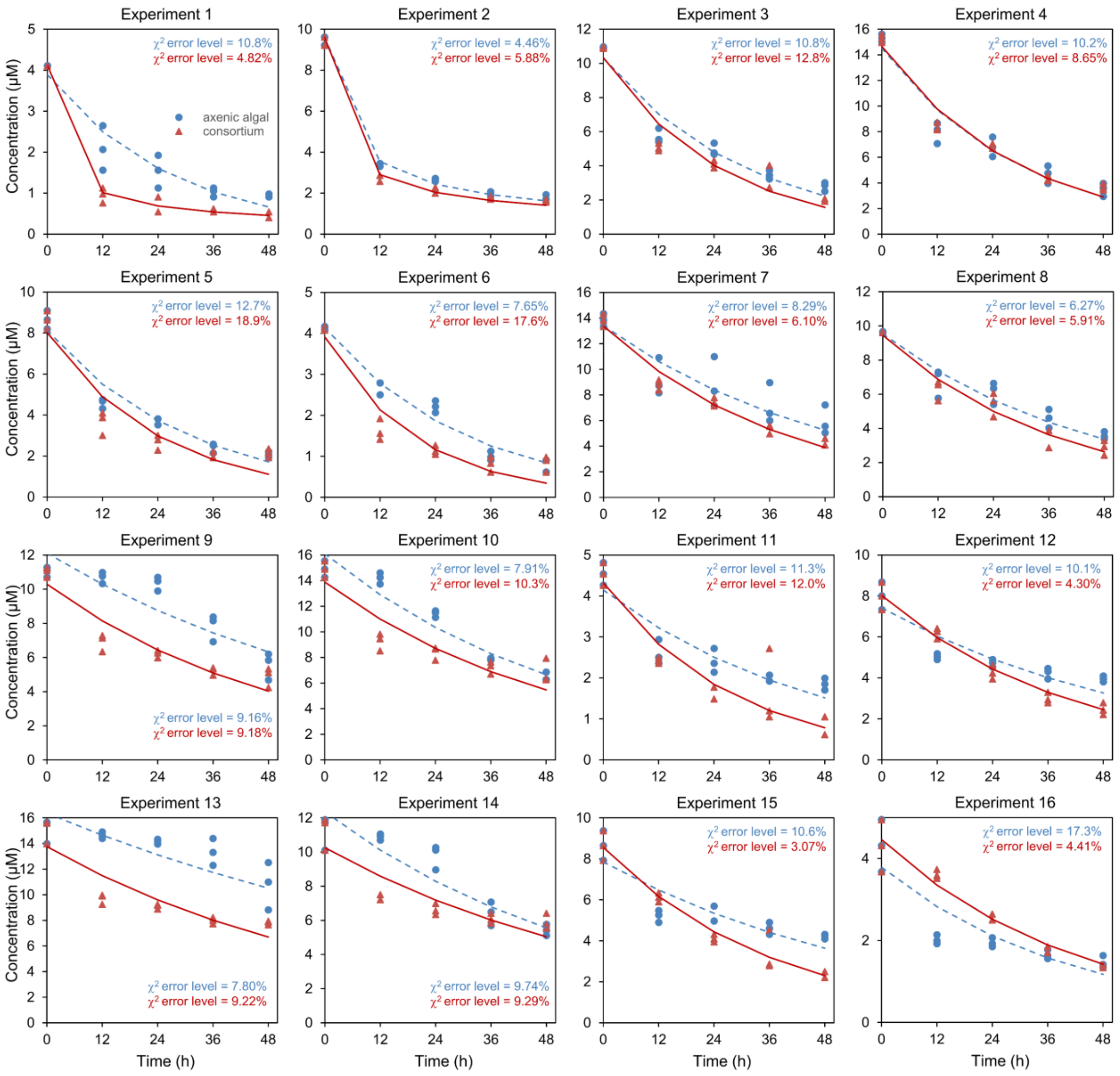


**Supplementary Fig. S5** Degradation of Acid Black 1 (AB1) dye under different experimental conditions of $L_{16}$ orthogonal array design (Experiment 1-16) in axenic algal alone (dotted blue lines) and algal-bacterial consortium (solid red lines) setups over a period of 48 hours.

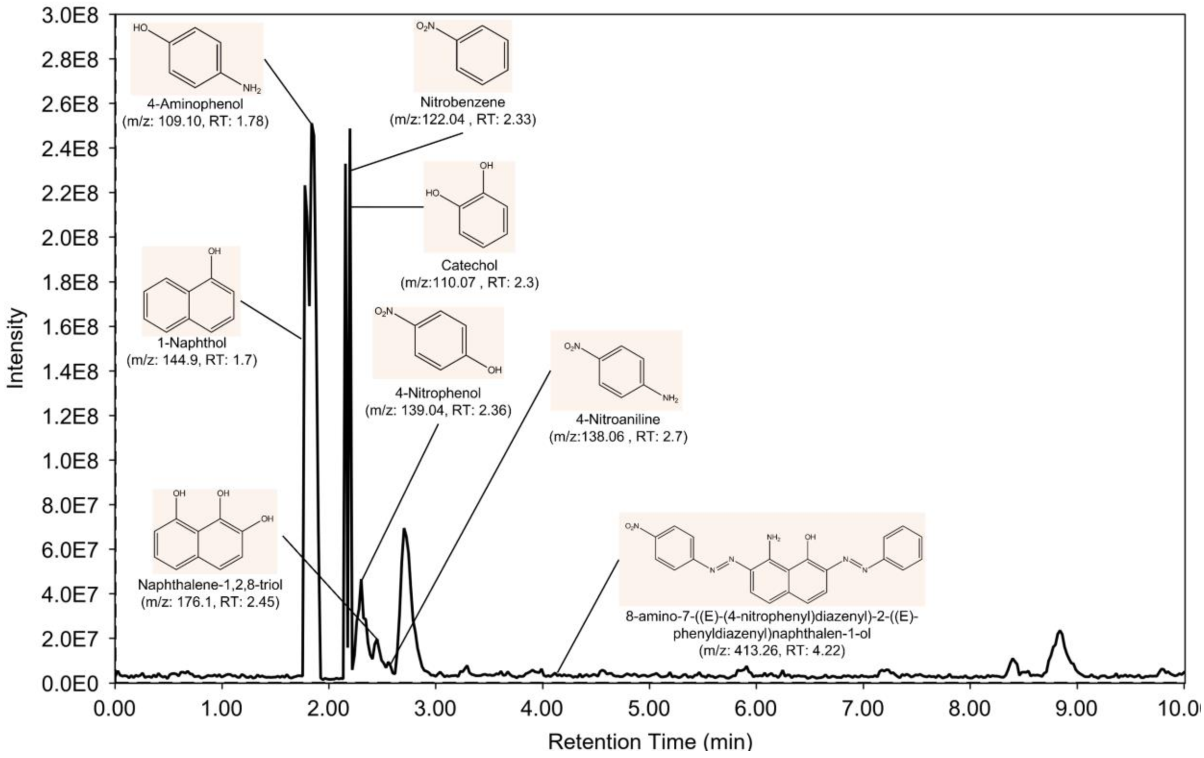


**Supplementary Fig. S6** The chromatograms of AB1 biodegraded products after LC-MS analysis.

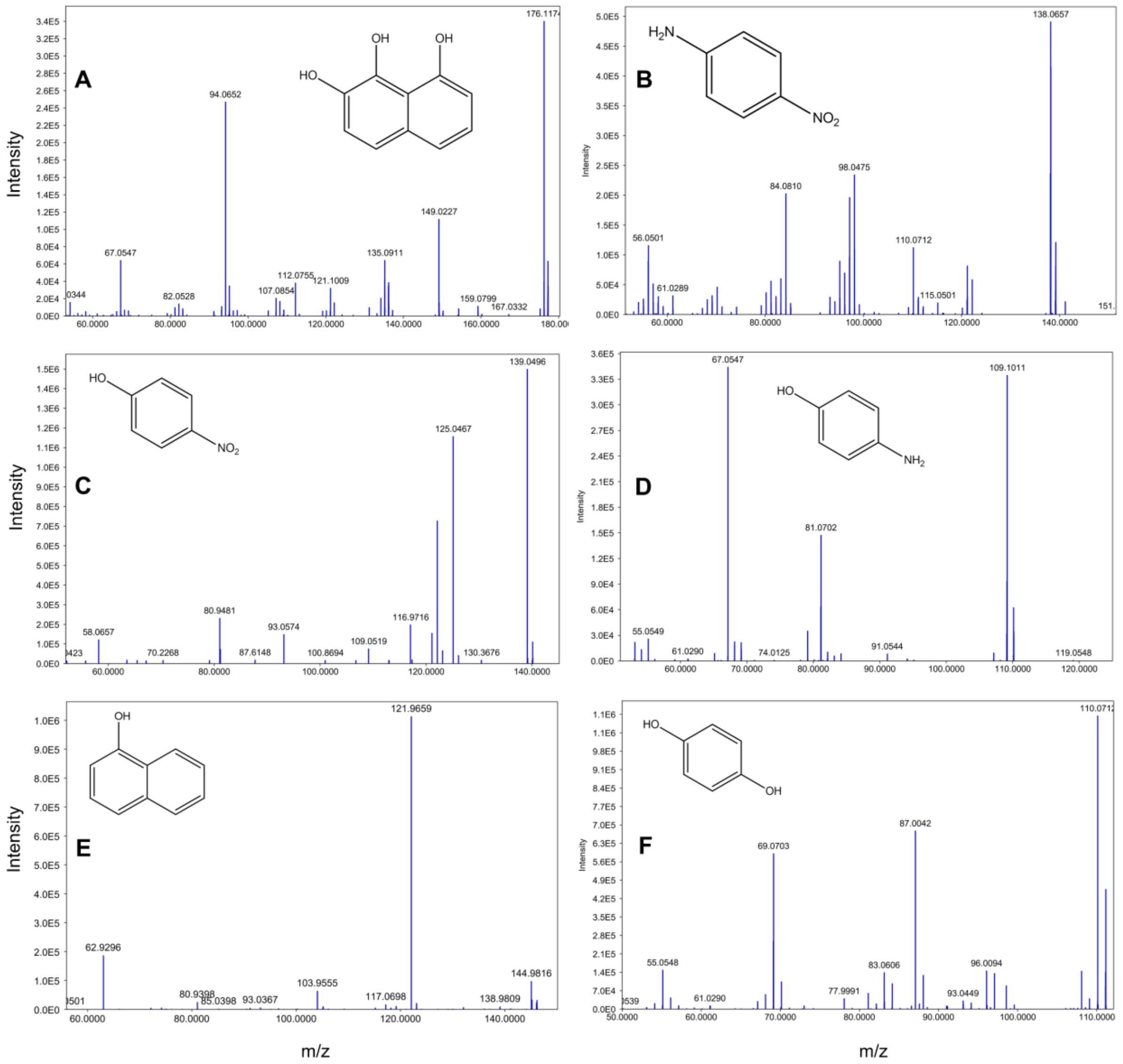


**Supplementary Fig. S7** Mass spectra of ABI biodegraded products showing formation of **A** Naphthalene-1,2,8-triol, **B** 4-Nitroaniline, **C** 4-Nitrophenol, **D** 4-Aminophenol, **E** Naphthalene-1-ol, and **F** Catechol.

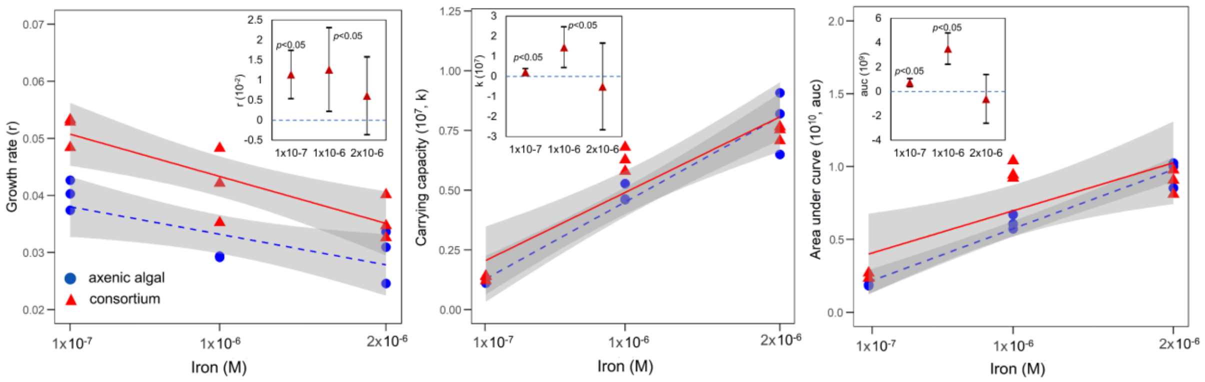


**Supplementary Fig. S8:** The confidence interval plots of different growth parameters of algal growth in consortium w.r.t. axenic growth (horizontal blue dashed line) suggests growth promotional properties of *Ralstonia pickettii* PW2 under low concentration of iron. The bacterium significantly enhanced the growth rate, carrying capacity, and area under curve of *Chlorella sorokiniana* at Fe concentration of $1x10^{-7}$ and $1x10^{-6}$ M. On the contrary, the effect of the presence of bacteria at a higher Fe concentration of $2x10^{-6}$ M was not significant (inset).

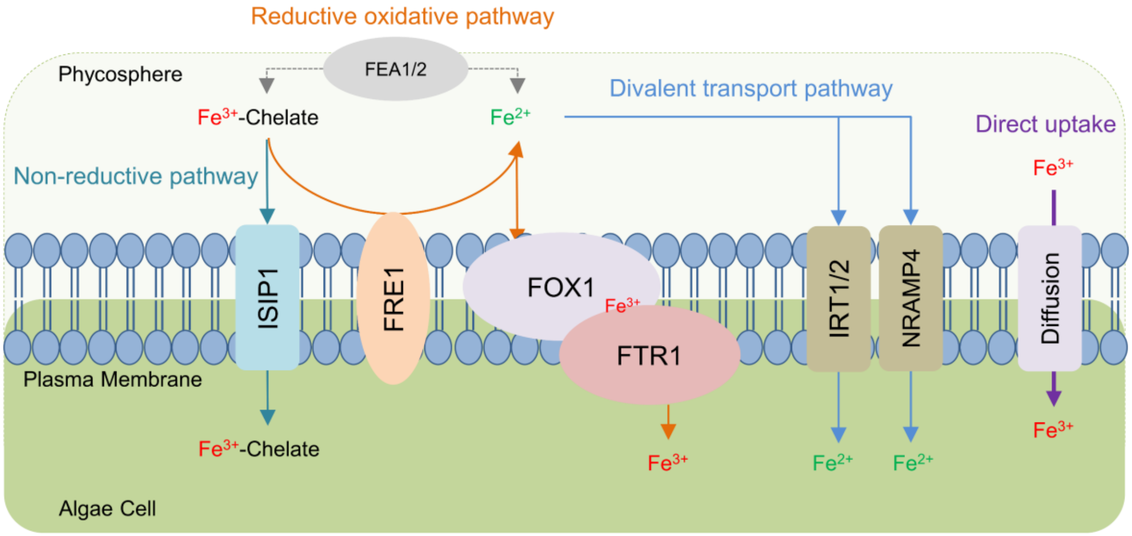


**Supplementary Fig. S9** The Fe-uptake mechanism in microalgae (based on the iron transport pathway deduced in green microalgae *Chlamydomonas reinhardtii* which as high similarity with *Chorella sorokininan*; Supplementary Data 4k-m) can be divided into the reductive oxidative pathway, divalent transport pathway, non-reductive uptake, and direct diffusion. The reductive oxidative pathway begins with the cell surface reduction of chelated $Fe^{3+}$ to $Fe^{2+}$ by a plasma membrane-bound NADPH-oxidoreductases (NOX family) FRE1 protein [2]. The extracellular FEA1 and FEA2 protein here plays a vital role in the accumulation of Fe-chelates near the cell surface as they bind with the ferric ($Fe^{3+}$) iron and make it available for reduction by FRE1 [3]. The second stage includes the further oxidation of ferrous ($Fe^{2+}$) iron by a multicopper ferroxidase (FOX1) and the transfer of $Fe^{3+}$ to a transporter ferric permease protein (FTR1). Allen et al., 2007 also postulated that FEA1/2 may also play a vital role in increasing the bioavailability of $Fe^{2+}$ for FOX1 as the proteins have been known to increase the solubility of ferrous iron. On the other hand, in a divalent transport pathway, the reduced $Fe^{2+}$ is directly incorporated into the cell via zip family proteins (IRT1 and IRT2) and natural resistance-associated macrophage proteins (NRAMP4). The zip family proteins have been known to transport divalent ions like zinc and iron into the cell. The direct incorporation of Fe-siderophore chelate has also been reported in an endocytosis-mediated uptake by iron starvation-induced protein 1 (ISIP1), which is a non-reductive iron uptake pathway [4]. Also, at higher iron concentration, it has been reported than ferric iron can diffuse directly via cell membrane, and gets further reduced inside the cell [5].

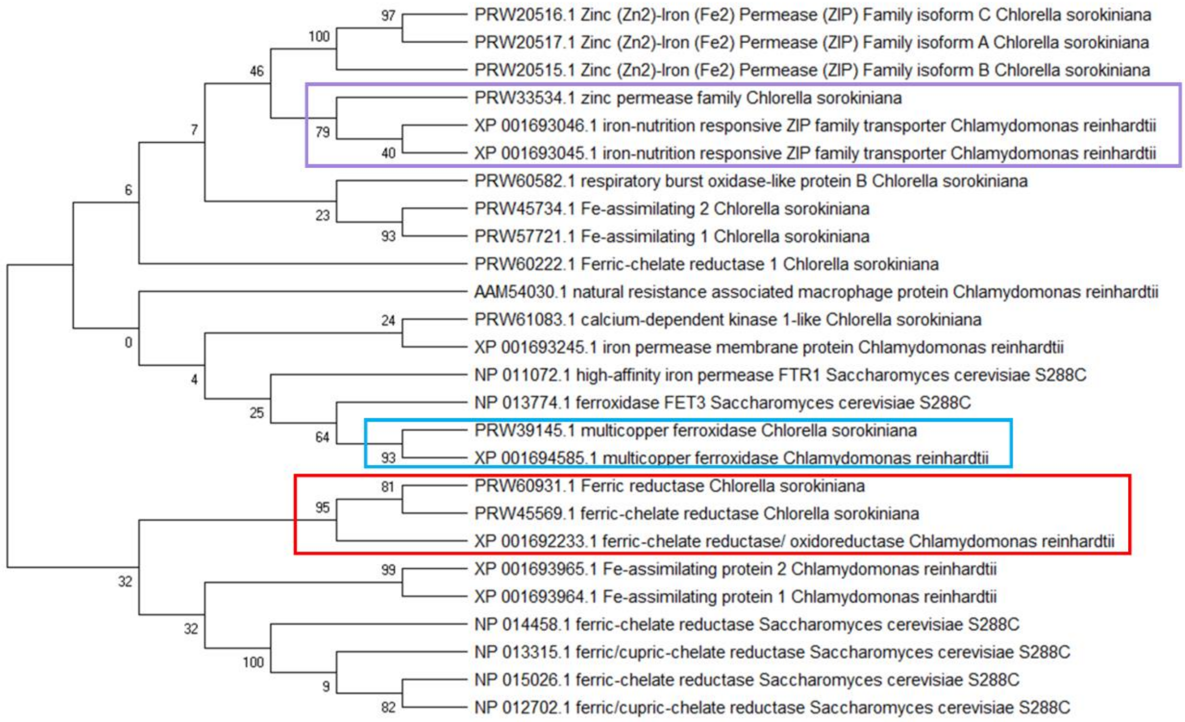


**Supplementary Fig. S10** The evolutionary history was inferred by using the Maximum Likelihood method and JTT matrix-based model in MEGA X software. The percentage of trees in which the associated taxa clustered together is shown next to the branches (a total of 500 bootstrap replication were used). This analysis involved 26 amino acid sequences from species *Chlorella sorokinana* UTEX 1602, *Chlamydomonas reinhardtii* and *Saccharomyces cerevisiae*. The protein sequences with maximum likelihood and higher bootstrap value are highlighted using colour codes. The red colour indicates the FRE1 (ferric- chelate reductase like) proteins showing a close relation between *Chlorella sorokiniana* and *Chlamydomonas reinhardtii*.

#### Supplementary Tables

**Supplementary Table 1:** The difference in the growth parameters of the microalgae *Chlorella sorokiniana* and *Scenedesmus* spp. co-cultured in the presence of siderophore producing bacteria.

| **Algae** | **Growth parameter** | **Co-culture treatments** | | | |
|---|---|---|---|---|---|
| | | axenic algal | *Serratia plymuthica* PW1 | *Ralstonia pickettii* PW2 | *Serratia liquefaciens* PW71 |
| *Chlorella sorokiniana* | Carrying capacity (k) | $4.82 \pm 0.45 \times 10^6$ | $3.34 \pm 1.99 \times 10^6$ | $4.97 \pm 0.13 \times 10^6$ | $3.40 \pm 0.21 \times 10^6$ |
| | Growth rate (r) | $1.95 \pm 0.33 \times 10^{-2}$ | $0.97 \pm 0.41 \times 10^{-2}$ | $5.02 \pm 1.09 \times 10^{-2}$ | $4.14 \pm 1.13 \times 10^{-2}$ |
| | Doubling time (Dt) | 35.45±3.905 | 70.98±10.29 | 13.80±1.060 | 16.71±0.4562 |
| | Area under curve (auc) | $6.01 \pm 0.49 \times 10^8$ | $3.26 \pm 1.51 \times 10^8$ | $11.01 \pm 0.37 \times 10^8$ | $7.05 \pm 0.25 \times 10^8$ |
| *Scenedesmus* spp. | Carrying capacity (k) | $11.25 \pm 0.83 \times 10^6$ | $5.99 \pm 0.27 \times 10^6$ | $9.85 \pm 0.38 \times 10^6$ | $9.88 \pm 0.66 \times 10^6$ |
| | Growth rate (r) | $2.60 \pm 0.51 \times 10^{-2}$ | $3.89 \pm 0.74 \times 10^{-2}$ | $2.63 \pm 0.27 \times 10^{-2}$ | $2.14 \pm 0.31 \times 10^{-2}$ |
| | Doubling time (Dt) | 26.65±5.337 | 17.79±20.78 | 26.28±5.641 | 32.36±6.411 |
| | Area under curve (auc) | $18.05 \pm 4.73 \times 10^8$ | $12.13 \pm 3.08 \times 10^8$ | $15.83 \pm 1.25 \times 10^8$ | $15.07 \pm 2.40 \times 10^8$ |

**Supplementary Table 2:** The output of Levene's, One-way ANOVA, and Tukey's posthoc tests used to compute the difference in the growth parameters of *Chlorella sorokiniana* and *Scenedesmus* spp. algae co-cultured in the presence of siderophore producing bacterial strains.

| Growth parameters | Algae | Levene | | ANOVA | | | Tukey | |
|---|---|---|---|---|---|---|---|---|
| | | Levene Statistic | *p* value | Sum of squares | F statistics | *p* value | Association of axenic algal w.r.t | *p* value |
| Growth rate (r) | *Chlorella sorokiniana* | 2.495 | 0.134 | 0.003 (Between groups) 0.000 (within groups) | 51.5 | 0.000 | consortium_PW1 | 0.068 |
| | | | | | | | consortium_PW2 | 0.000 |
| | | | | | | | consortium_PW71 | 0.002 |
| | *Scenedesmus* spp. | 7.858 | 0.009 | 0.005 (Between groups) 0.005 (within groups) | 11.731 | 2.632 | consortium_PW1 | 0.261 |
| | | | | | | | consortium_PW2 | 0.989 |
| | | | | | | | consortium_PW71 | 0.968 |
| Carrying capacity (k) | *Chlorella sorokiniana* | 1.707 | 0.242 | $8.77x10^{12}$ (Between groups) $3.50x10^{12}$ (within groups) | 6.674 | 0.014 | consortium_PW1 | 0.004 |
| | | | | | | | consortium_PW2 | 1.000 |
| | | | | | | | consortium_PW71 | 0.067 |
| | *Scenedesmus* spp. | 3.726 | 0.061 | $4.91x10^{13}$ (Between groups) $4.80x10^{12}$ (within groups) | 27.273 | 0.000 | consortium_PW1 | 0.000 |
| | | | | | | | consortium_PW2 | 0.231 |
| | | | | | | | consortium_PW71 | 0.263 |
| Doubling time (Dt) | *Chlorella sorokiniana* | 4.468 | 0.04 | 5690.06 (Between groups) 737.36 (within groups) | 10.362 | 20.578 | consortium_PW1 | 0.009 |
| | | | | | | | consortium_PW2 | 0.145 |
| | | | | | | | consortium_PW71 | 0.218 |
| | *Scenedesmus* spp. | 0.352 | 0.789 | 623.2 (Between groups) 325.73 (within groups) | 5.102 | 0.029 | consortium_PW1 | 0.261 |
| | | | | | | | consortium_PW2 | 0.788 |
| | | | | | | | consortium_PW71 | 0.391 |
| Area under curve (auc) | *Chlorella sorokiniana* | 2.564 | 0.128 | $8.92x10^{17}$ (Between groups) $2.75x10^{16}$ (within groups) | 86.391 | 0.000 | consortium_PW1 | 0.001 |
| | | | | | | | consortium_PW2 | 0.000 |
| | | | | | | | consortium_PW71 | 0.430 |
| | *Scenedesmus* spp. | 4.66 | 0.036 | $6.24x10^{17}$ (Between groups) $1.75x10^{17}$ (within groups) | 9.513 | 0.005 | consortium_PW1 | 0.003 |
| | | | | | | | consortium_PW2 | 0.086 |
| | | | | | | | consortium_PW71 | 0.033 |

**Supplementary Table 3:** The output of the AB1 degradation kinetic modeling performed using '*mkin*' package in R.

| Treatment setup | Kinetic model | Chi-square Error (%) | Half-life (h) | Degradation after 144 h (%) |
|---|---|---|---|---|
| axenic algal | SFO | 6.015 | 120 ± 9.43 | 54.40 ± 2.28 |
| consortium | SFO | 8.557 | 65.40 ± 0.50 | 73.96 ± 0.28 |
| axenic bacterial | NA | NA | NA | 8.15 ± 0.32 |
| axenic algal (EDTA) | FOMC | 4.202 | 21.59 ± 1.89 | 86.21 ± 0.85 |
| consortium (EDTA) | FOMC | 2.859 | 15.27 ± 2.49 | 89.99 ± 0.05 |
| axenic bacterial (EDTA) | NA | NA | NA | 6.16 ± 0.24 |

**Supplementary Table 4:** The output of Levene's, One-way ANOVA, and Tukey's posthoc tests used to compute the difference in the half-life of AB1 in axenic algal and algal-bacterial consortium treatment setups with varied conditions of iron bioavailability.

| Parameter | Levene | | ANOVA | | | Tukey | | | |
|---|---|---|---|---|---|---|---|---|---|
| | Levene Statistic | *p* value | Sum of squares | F statistics | *p* value | Association of axenic algal w.r.t. | *p* value | Association of axenic algal (EDTA) w.r.t. | *p* value |
| Half-life | 4.006 | 0.052 | 21079.95 (Between groups) 593.679 (within groups) | 94.686 | 0.000 | consortium | 0.000 | consortium | 0.000 |
| | | | | | | axenic algal (EDTA) | 0.000 | axenic algal | 0.001 |
| | | | | | | consortium (EDTA) | 0.000 | consortium (EDTA) | 0.807 |

**Supplementary Table 5:** The L'16 ($4^{(3)}$) orthogonal array design with three factors (Fe conc., pH, and Dye conc.) with their respective levels. The responses were recorded as the rate of degradation ($h^{-1}$) of Acid Black 1 (AB1) dye in the microbial setups with only axenic algal (setup 1) and algal-bacterial consortium (setup 2).

| Exp. | Factors | | | Response (Rate, $h^{-1}$) | | Degradation (%) | |
|---|---|---|---|---|---|---|---|
| | Fe conc. | pH | Dye conc. | axenic algal (setup 1) | consortium (setup 2) | axenic algal (setup 1) | consortium (setup 2) |
| 1 | $1x10^{-7}$ | pH6 | 4 µM | $3.69 \pm 0.55$ x $10^{-2}$ | $9.10 \pm 1.23$ x $10^{-2}$ | 77.36 ± 0.58 | 89.12 ± 1.17 |
| 2 | $1x10^{-7}$ | pH7 | 8 µM | $5.34 \pm 0.19$ x $10^{-2}$ | $6.73 \pm 0.09$ x $10^{-2}$ | 81.53 ± 0.75 | 83.04 ± 0.43 |
| 3 | $1x10^{-7}$ | pH8 | 12 µM | $3.19 \pm 0.13$ x $10^{-2}$ | $3.93 \pm 0.17$ x $10^{-2}$ | 74.45 ± 1.38 | 81.53 ± 0.44 |
| 4 | $1x10^{-7}$ | pH9 | 16 µM | $3.39 \pm 0.21$ x $10^{-2}$ | $3.38 \pm 0.14$ x $10^{-2}$ | 77.17 ± 1.94 | 76.07 ± 0.98 |
| 5 | $1x10^{-6}$ | pH6 | 8 µM | $3.24 \pm 0.08$ x $10^{-2}$ | $4.21 \pm 0.45$ x $10^{-2}$ | 76.10± 0.96 | 75.26 ± 1.45 |
| 6 | $1x10^{-6}$ | pH7 | 4 µM | $3.32 \pm 0.21$ x $10^{-2}$ | $5.14 \pm 0.56$ x $10^{-2}$ | 80.37 ± 2.34 | 79.78 ± 2.68 |
| 7 | $1x10^{-6}$ | pH8 | 16 µM | $1.97 \pm 0.15$ x $10^{-2}$ | $2.57 \pm 0.08$ x $10^{-2}$ | 57.15 ± 4.74 | 67.97 ± 1.22 |
| 8 | $1x10^{-6}$ | pH9 | 12 µM | $2.17 \pm 0.06$ x $10^{-2}$ | $2.66 \pm 0.13$ x $10^{-2}$ | 63.01 ± 1.32 | 70.03 ± 2.61 |
| 9 | $2x10^{-6}$ | pH6 | 12 µM | $1.36 \pm 0.11$ x $10^{-2}$ | $1.95 \pm 0.12$ x $10^{-2}$ | 49.50 ± 4.16 | 55.64 ± 3.01 |
| 10 | $2x10^{-6}$ | pH7 | 16 µM | $1.84 \pm 0.06$ x $10^{-2}$ | $1.95 \pm 0.11$ x $10^{-2}$ | 56.59 ± 1.29 | 54.00 ± 3.65 |
| 11 | $2x10^{-6}$ | pH8 | 4 µM | $2.08 \pm 0.06$ x $10^{-2}$ | $3.56 \pm 0.18$ x $10^{-2}$ | 59.26 ± 1.84 | 80.03 ± 3.19 |
| 12 | $2x10^{-6}$ | pH9 | 8 µM | $1.70 \pm 0.08$ x $10^{-2}$ | $2.47 \pm 0.18$ x $10^{-2}$ | 50.65 ± 1.04 | 69.06 ± 2.11 |
| 13 | $5x10^{-6}$ | pH6 | 16 µM | $1.06 \pm 0.09$ x $10^{-2}$ | $1.50 \pm 0.10$ x $10^{-2}$ | 28.58 ± 7.11 | 50.06 ± 0.57 |
| 14 | $5x10^{-6}$ | pH7 | 12 µM | $1.67 \pm 0.12$ x $10^{-2}$ | $1.48 \pm 0.04$ x $10^{-2}$ | 51.73 ± 1.67 | 47.43 ± 2.32 |
| 15 | $5x10^{-6}$ | pH8 | 8 µM | $1.61 \pm 0.14$ x $10^{-2}$ | $2.74 \pm 0.21$ x $10^{-2}$ | 51.24 ± 0.73 | 72.19 ± 1.11 |
| 16 | $5x10^{-6}$ | pH9 | 4 µM | $2.48 \pm 0.25$ x $10^{-2}$ | $2.39 \pm 0.20$ x $10^{-2}$ | 60.22 ± 2.01 | 68.34 ± 0.56 |

**Supplementary Table 6:** The output of the multiple linear regression model used to compute the difference in the rate of AB1 degradation in the 16 experiments suggested by Taguchi's $L_{16}$ $(4^3)$ orthogonal array in two different setups.

| Treatments | Factors | Rank (delta) | Multiple linear regression equation | $R^2$ | F statistics | *p* value |
|---|---|---|---|---|---|---|
| axenic algal (setup 1) | Fe conc. | 1 (0.022) | Rate = 0.0252 + 0.014 ($1x10^{-7}$) + 0.001 ($1x10^{-6}$) – 0.007 ($2x10^{-6}$) | 94.81% | 27.12 | 0.001 |
| | pH | 3 (0.008) | Rate = 0.0252 - 0.001 (pH6) + 0.005 (pH7) – 0.003 (pH8) | | 3.2 | 0.105 |
| | Dye conc. | 2 (0.009) | Rate = 0.0252 + 0.004 (4µM) + 0.004 (8µM) – 0.004 (12µM) | | 6.25 | 0.028 |
| consortium (setup 2) | Fe conc. | 1 (0.03) | Rate = 0.0348 + 0.022 ($1x10^{-7}$) + 0.001 ($1x10^{-6}$) – 0.01 ($2x10^{-6}$) | 94.15% | 18.46 | 0.002 |
| | pH | 3 (0.01) | Rate = 0.0348 - 0.007 (pH6) + 0.003 (pH7) – 0.003 (pH8) | | 2.78 | 0.133 |
| | Dye conc. | 2 (0.02) | Rate = 0.0348 + 0.015 (4µM) + 0.005 (8µM) – 0.009 (12µM) | | 10.92 | 0.008 |

**Supplementary Table 7:** Estimated model coefficients for means of the 16 experiments suggested by Taguchi's $L_{16}$ $(4^3)$ orthogonal array in two different setups.

| Factors | axenic algal (setup 1) | | | consortium (setup 2) | | |
|---|---|---|---|---|---|---|
| | Coefficient | t Statistics | *p* value | Coefficient | t Statistics | *p* value |
| Constant | 0.025231 | 24.927 | 0 | 0.034893 | 17.881 | 0 |
| Fe conc. ($1x10^{-7}$) | 0.014302 | 8.158 | 0 | 0.022976 | 6.798 | 0 |
| Fe conc. ($1x10^{-6}$) | 0.001589 | 0.906 | 0.4 | 0.001622 | 0.48 | 0.648 |
| Fe conc. ($2x10^{-6}$) | -0.00773 | -4.409 | 0.005 | -0.01002 | -2.963 | 0.025 |
| pH (pH6) | -0.00136 | -0.776 | 0.467 | 0.007054 | 2.087 | 0.082 |
| pH (pH7) | 0.005244 | 2.991 | 0.024 | 0.003392 | 1.004 | 0.354 |
| pH (pH8) | -0.00305 | -1.737 | 0.133 | -0.00286 | -0.846 | 0.43 |
| Dye conc. (4µM) | 0.004215 | 2.404 | 0.053 | 0.015617 | 4.62 | 0.004 |
| Dye conc. (8µM) | 0.004547 | 2.594 | 0.041 | 0.005527 | 1.635 | 0.153 |
| Dye conc. (12µM) | -0.00422 | -2.409 | 0.053 | -0.0098 | -2.899 | 0.027 |

**Supplementary Table 8:** The difference in the growth parameters of the microalgae *Chlorella sorokiniana* co-cultured in the presence of siderophore producing bacteria *Ralstonia pickettii* PW2 at different concentrations of iron.

| Growth parameter | $1x10^{-7}$ M | | $1x10^{-6}$ M | | $2x10^{-6}$ M | |
|---|---|---|---|---|---|---|
| | axenic algal | consortium | axenic algal | consortium | axenic algal | consortium |
| Carrying capacity (k) | 1.11±0.007 $x10^7$ | 1.37±0.06 $x10^7$ | 4.83±0.22 $x10^7$ | 6.28±0.29 $x10^7$ | 7.91±0.75 $x10^7$ | 7.41±0.17 $x10^7$ |
| Growth rate (r) | 4.01±0.15 $x10^{-2}$ | 5.15±0.15 $x10^{-2}$ | 2.92±0.006 $x10^{-2}$ | 4.18±0.37 $x10^{-2}$ | 2.97±0.26 $x10^{-2}$ | 3.58±0.22 $x10^{-2}$ |
| Doubling time (Dt) | 17.32±0.66 | 13.48±0.42 | 23.70±0.05 | 16.82±1.54 | 23.73±2.29 | 19.50±1.17 |
| Area under curve (auc) | 1.86±0.02 $x10^9$ | 2.57±0.11 $x10^9$ | 6.17±0.28 $x10^9$ | 9.68±0.36 $x10^9$ | 9.59±0.53 $x10^9$ | 8.98±0.48 $x10^9$ |

**Supplementary Table 9:** The output of the linear regression model used to compute the difference in the growth parameters of *Chlorella sorokiniana* co-cultured (with and without bacteria) at the varying concentrations of iron.

| Growth parameters | Fe conc. | Linear regression equation | $R^2$ | F statistics | t value | *p* value |
|---|---|---|---|---|---|---|
| Growth rate (r) | $1x10^{-7}$ | r = 0.0401 + 0.011 consortium | 87.23% | 27.33 | 5.228 | 0.006 |
| | $1x10^{-6}$ | r = 0.029 + 0.012 consortium | 73.82% | 11.28 | 3.358 | 0.028 |
| | $2x10^{-6}$ | r = 0.029 + 0.006 consortium | 42.99% | 3.07 | 1.737 | 0.157 |
| Carrying capacity (k) | $1x10^{-7}$ | k = $1.1x10^7$ + $0.2x10^7$ consortium | 75.17% | 12.11 | 3.48 | 0.025 |
| | $1x10^{-6}$ | k = $4.8x10^7$ + $1.4x10^7$ consortium | 79.76% | 15.77 | 3.971 | 0.0165 |
| | $2x10^{-6}$ | k = 7.9 $x10^7$ – $0.5x10^7$ consortium | 9.53% | 0.42 | -0.642 | 0.551 |
| Area under curve (auc) | $1x10^{-7}$ | auc = $1.8x10^9$ + $0.7x10^9$ consortium | 90.17% | 36.68 | 6.052 | 0.003 |
| | $1x10^{-6}$ | auc = $6.1x10^9$ + $3.5x10^9$ consortium | 93.44% | 56.97 | 7.548 | 0.001 |
| | $2x10^{-6}$ | auc = 9.5 $x10^9$ - 0.6 $x10^9$ consortium | 15.22% | 0.718 | -0.847 | 0.444 |

**Supplementary Table 10:** The output of the linear regression model used to compute the difference in the ferrireductase activity of *Chlorella sorokiniana* co-cultured (with and without bacteria) at the varying concentrations of iron.

| Fe conc. | Linear regression equation | $R^2$ | F statistics | t value | *p* value |
|---|---|---|---|---|---|
| $1x10^{-7}$ | Ferri = 0.015 + 0.009 consortium | 94.70% | 71.51 | 8.456 | 0.001 |
| $1x10^{-6}$ | Ferri = 0.006 + 0.002 consortium | 89.92% | 35.69 | 5.974 | 0.003 |
| $2x10^{-6}$ | Ferri = 0.004 – 0.0007 consortium | 19.71% | 0.98 | 1.737 | 0.377 |

**Supplementary Table 11:** The output of the linear regression model used to compute the difference in the ferrireductase activity of *Chlorella sorokiniana* cultured at the varying concentrations of ferrireductase inhibitor DPI.

| S.No. | DPI conc. | Linear regression equation | $R^2$ | F statistics | t value | *p* value |
|---|---|---|---|---|---|---|
| 1 | 50µM | Ferri = 0.023 + 0.009 50µM | 98.31% | 155.4 | -18.74 | 0.000 |
| 2 | 100µM | Ferri = 0.023 + 0.002 100µM | | | -18.51 | 0.000 |
| 3 | 150µM | Ferri = 0.023 - 0.0007 150µM | | | -13.97 | 0.000 |

**Supplementary Table 12:** The output of Levene's, One-way ANOVA, and Tukey's posthoc tests used to compute the difference in the azoreductase activity of *Chlorella sorokiniana* cultured with and without DPI and iron.

| Assay | Levene | | ANOVA | | | Tukey | |
|---|---|---|---|---|---|---|---|
| | **Levene Statistic** | ***p* value** | **Sum of squares** | **F statistics** | ***p* value** | **Association wrt DPI-Fe-** | ***p* value** |
| Azoreductase | 0.0482 | 0.704 | 0.000 (Between groups) 0.000 (within groups) | 31.692 | 0.000 | DPI-Fe+ | 0.012 |
| | | | | | | DPI+Fe- | 0.035 |
| | | | | | | DPI+Fe+ | 0.008 |

**Supplementary Table 13:** The similarities between Ferric reductase and iron transporter proteins in green algae *Chlamydomonas reinhardtii* and *Chlorella sorokiniana*

| S.No. | *Chlamydomonas reinhardtii* | *Chlorella sorokiniana* UTEX 1602 | Similarity |
|---|---|---|---|
| 1 | Ferric-chelate reductase/ oxidoreductase (FRE1) https://www.ncbi.nlm.nih.gov/protein/XP_001692233.1 | Ferric-chelate reductase https://www.ncbi.nlm.nih.gov/protein/PRW45569.1 | High |
| | | Ferric reductase https://www.ncbi.nlm.nih.gov/protein/PRW60931.1 | High |
| | | Ferric-chelate reductase 1 https://www.ncbi.nlm.nih.gov/protein/PRW60222.1 | Low |
| 2 | Fe-assimilating protein 1 and 2 https://www.ncbi.nlm.nih.gov/protein/XP_001693964.1 https://www.ncbi.nlm.nih.gov/protein/XP_001693965.1 | Fe-assimilating 1/ 2 https://www.ncbi.nlm.nih.gov/protein/PRW57721.1/ https://www.ncbi.nlm.nih.gov/protein/PRW45734.1/ | Low |
| 3 | Multicopper ferrioxidase (FOX1) https://www.ncbi.nlm.nih.gov/protein/XP_001694585.1 | Multicopper ferroxidase https://www.ncbi.nlm.nih.gov/protein/PRW39145.1 | High |
| 4 | Iron permease, membrane protein (FTR1) https://www.ncbi.nlm.nih.gov/protein/XP_001693245.1 | Calcium-dependent kinase 1-like https://www.ncbi.nlm.nih.gov/protein/PRW61083.1 | Low |
| 5 | Iron-nutrition responsive ZIP family transporter (IRT1/2) https://www.ncbi.nlm.nih.gov/protein/XP_001693045.1 | Zinc permease family https://www.ncbi.nlm.nih.gov/protein/PRW33534.1 | High |

**Supplementary Table 14:** List of genes and enzymes in the plasma membrane of algae and yeast associated with iron transport. The proteins in *Chlorella sorokiniana* with similar functions as those of *Chlamydomonas reinhardtii* and *Saccharomyces cerevisiae* were identified by using the NCBI BLAST.

| Species | Gene | Protein and function | Uniprot/NCBI Link | Reference |
|---|---|---|---|---|
| **Green Algae-** *Chlamydomonas reinhardtii* | *FRE1* | **Ferric-chelate reductase/ oxidoreductase:** Cell surface iron reductases that reduce Ferric-citrate and siderophore-bound iron | Uniprot: https://www.uniprot.org/uniprot/A2I2U7 NCBI: https://www.ncbi.nlm.nih.gov/protein/XP_001692233.1 | [3, 6] |
| | *FOX1* | **Multicopper ferrioxidase:** Plasma membrane-associated multicopper oxidase that reconverts Fe(II) to Fe(III) which is a substrate for associated trivalent cation—specific permease FTR1 | Uniprot: https://www.uniprot.org/uniprot/A8IZT9 NCBI: https://www.ncbi.nlm.nih.gov/protein/XP_001694585.1 | |
| | *FTR1* | **Iron permease, membrane protein:** Ferric permease transports Fe(III) into the cell | Uniprot: https://www.uniprot.org/uniprot/Q8LL16 NCBI: https://www.ncbi.nlm.nih.gov/protein/XP_001693245.1 | |
| | *FEA1 / FEA2* | **Fe-assimilating protein 1 and 2:** Responds to iron deficiency in assimilation of iron for FRE1 and FOX1 | Uniprot: https://www.uniprot.org/uniprot/Q9LD42 https://www.uniprot.org/uniprot/Q38J95 NCBI: https://www.ncbi.nlm.nih.gov/protein/XP_001693964.1 https://www.ncbi.nlm.nih.gov/protein/XP_001693965.1 | |
| | *IRT1/2* | **Iron-nutrition responsive ZIP family transporter:** Transportation of divalent metals ions through plasma membrane | Uniprot: https://www.uniprot.org/uniprot/A8IW06 https://www.uniprot.org/uniprot/A8IW07 NCBI: https://www.ncbi.nlm.nih.gov/protein/XP_001693045.1 https://www.ncbi.nlm.nih.gov/protein/XP_001693046.1 | |
| | *NRAMP4* | **Natural resistance-associated macrophage protein 4:** Transportation of divalent metals ions through plasma membrane | NCBI: https://www.ncbi.nlm.nih.gov/protein/AAM54030.1 | [6] |
| **Baker's Yeast-** *Saccharomyces cerevisiae* (strain ATCC 204508 / S288c) | *FRE1/2/3/4* | **Ferric/cupric reductase transmembrane component 1 and 2:** Cell surface iron reductases that reduce Ferric-citrate and siderophore-bound iron **Ferric/cupric reductase transmembrane component 3:** Selectivity for iron in complex with hydroxamate-type siderophore **Ferric/cupric reductase transmembrane component 4:** | Uniprot: https://www.uniprot.org/uniprot/P32791 https://www.uniprot.org/uniprot/P36033 https://www.uniprot.org/uniprot/Q08905 https://www.uniprot.org/uniprot/P53746 NCBI: https://www.ncbi.nlm.nih.gov/protein/NP_013315.1 https://www.ncbi.nlm.nih.gov/protein/NP_012702.1 https://www.ncbi.nlm.nih.gov/protein/NP_015026.1 https://www.ncbi.nlm.nih.gov/protein/NP_014458.1 | [3, 7, 8] |

| | | | | |
|---|---|---|---|---|
| | | Low-affinity activity on rhodotorulic acid (hydroxamate siderophore) | | |
| | *FET3* | **Iron transport multicopper oxidase FET3/ferroxidaseFET3:** Plasma membrane-associated multicopper oxidase that reconverts Fe(II) to Fe(III) which is a substrate for associated trivalent cation specific Ftr1p | Uniprot: https://www.uniprot.org/uniprot/P38993 NCBI: https://www.ncbi.nlm.nih.gov/protein/NP_013774.1 | |
| | *FTR1* | **Plasma membrane iron permease/ high-affinity iron permease FTR1 :** Ferric permease transports Fe(III) into the cell | Uniprot: https://www.uniprot.org/uniprot/P40088 NCBI: https://www.ncbi.nlm.nih.gov/protein/NP_011072.1 | |
| **Green algae-** *Chlorella sorokinana* UTEX 1602 | *C2E21_5734* | **Ferric-chelate reductase** | Uniprot: https://www.uniprot.org/uniprot/A0A2P6TMJ2 NCBI: https://www.ncbi.nlm.nih.gov/protein/PRW45569.1 | Maybe like FRE1 |
| | *C2E21_0396* | **Ferric reductase** | Uniprot: https://www.uniprot.org/uniprot/A0A2P6U3N8 NCBI: https://www.ncbi.nlm.nih.gov/protein/PRW60931.1 | |
| | *C2E21_1040* | **Ferric-chelate reductase 1** | Uniprot: https://www.uniprot.org/uniprot/A0A2P6U1N0 NCBI: https://www.ncbi.nlm.nih.gov/protein/PRW60222.1 | |
| | *C2E21_7043* | **Multicopper ferroxidase** | Uniprot: https://www.uniprot.org/uniprot/A0A2P6TIS5 NCBI: https://www.ncbi.nlm.nih.gov/protein/PRW39145.1 | Maybe like FOX1 and FET3 |
| | *C2E21_0141* | **Calcium-dependent kinase 1-like** | Uniprot: https://www.uniprot.org/uniprot/A0A2P6U456 NCBI: https://www.ncbi.nlm.nih.gov/protein/PRW61083.1 | Maybe like FTR1 |
| | *C2E21_3645/ C2E21_6014* | **Fe-assimilating 1/ 2** | Uniprot: https://www.uniprot.org/uniprot/A0A2P6TUJ1 https://www.uniprot.org/uniprot/A0A2P6TN26 NCBI: https://www.ncbi.nlm.nih.gov/protein/PRW57721.1/ https://www.ncbi.nlm.nih.gov/protein/PRW45734.1/ | Maybe like FEA1/2 |
| | *C2E21_8904* | **Zinc (Zn2)-Iron (Fe2) Permease (ZIP) Family isoform A, B, and C** | Uniprot: https://www.uniprot.org/uniprot/A0A2P6TCZ6 https://www.uniprot.org/uniprot/A0A2P6TCZ9 https://www.uniprot.org/uniprot/A0A2P6TD01 NCBI: https://www.ncbi.nlm.nih.gov/protein/PRW20517.1 https://www.ncbi.nlm.nih.gov/p | Maybe like IRT1/2 |

| | | | | |
|---|---|---|---|---|
| | | | rotein/PRW20515.1 https://www.ncbi.nlm.nih.gov/protein/PRW20516.1 | |
| | *C2E21_7513* | **Zinc permease family** | Uniprot: https://www.uniprot.org/uniprot/A0A2P6TGW7 NCBI: https://www.ncbi.nlm.nih.gov/protein/PRW33534.1 | Maybe like IRT1/2 |
| **Marine algae-** *Phaeodactylum tricornutum* 1055/1 | *ISIP1* | **Iron-starvation induced protein:** Uptake of siderophore bound iron directly into cell | Uniprot: https://www.uniprot.org/uniprot/B7GA90 | [4] |
| **Green alage-** *Chlorella vulgaris* | NA | Ferric reductase enzyme activity for iron uptake of hydroxymate siderophore bound iron. | NA | [9] |
| **Marine green algae-** *Chlorococcum littorale* | NA | Ferric reductase enzyme activity (BPDS-Fe(II) assay) | NA | [10] |
| **Green alage-** *Chlorella kessleri* | NA | Ferric chelate reductase enzyme activity (BPDS-Fe(II) assay) | NA | [11] |
| **Marine algae:** *Scrippsiella trochoidea* | NA | Ferric reductase enzyme activity (BPDS-Fe(II) assay) | NA | [12] |

**Supplementary Data References**


1. Sprouffske K, Wagner A. Growthcurver: An R package for obtaining interpretable metrics from microbial growth curves. *BMC Bioinformatics* 2016; **17**: 172.

2. Amin SA. The role of siderophores in algal-bacterial interactions in the marine environment. 2010. UC San Diego.
